# The histone chaperone Spt6 controls chromatin structure through its conserved N-terminal domain

**DOI:** 10.1101/2024.11.25.625227

**Authors:** James L. Warner, Vanda Lux, Václav Veverka, Fred Winston

## Abstract

The disassembly and reassembly of nucleosomes by histone chaperones is an essential activity during eukaryotic transcription elongation. This highly conserved process maintains chromatin integrity by transiently removing nucleosomes as barriers and then restoring them in the wake of transcription. While transcription elongation requires multiple histone chaperones, there is little understanding of how most of them function and why so many are required. Here, we show that the histone chaperone Spt6 acts through its acidic, intrinsically disordered N-terminal domain (NTD) to bind histones and control chromatin structure. The Spt6 NTD is essential for viability and its histone binding activity is conserved between yeast and humans. The essential nature of the Spt6 NTD can be bypassed by changes in another histone chaperone, FACT, revealing a close functional connection between the two. Our results have led to a mechanistic model for dynamic cooperation between multiple histone chaperones during transcription elongation.

## Introduction

During transcription elongation, RNA polymerase II (RNAPII) encounters a formidable barrier approximately every 200 base pairs – the nucleosome, the basic unit of chromatin. Nucleosomes consist of 147 bp of DNA wrapped around one H3/H4 tetramer and two H2A/H2B dimers^1^. As transcription proceeds, every nucleosome must be unwrapped, have its histones transferred from ahead of to behind RNAPII, and then be rewrapped. This process is essential for transcription to proceed and crucial to maintain chromatin integrity, as a reduced level of nucleosomes results in widespread changes in the fidelity of transcription initiation^2^.

RNAPII elongation through nucleosomes requires several histone chaperones^3–5^. Histone chaperones are a diverse set of histone-binding proteins that function in an ATP-independent fashion to allow transcription and other chromatin-related processes to occur^6,7^. Extensive biochemical and structural studies have provided an understanding of how one of these chaperones, FACT, interacts with histones during transcription elongation^8–12^. However, much less is understood about the other histone chaperones associated with RNAPII. It is unlikely that their functions are redundant, as at least four (Spt6, FACT, Spt5, and Spn1/IWS1) are individually essential for viability in both *S. cerevisiae* and humans^13–18^, demonstrating that each carries out a vital function. Furthermore, depletion of any of these factors causes widespread changes in transcription and chromatin structure^19–25^. Additional factors, including Elf1 and Spt2, although not essential for viability, are also important during elongation through nucleosomes^8,26–29^. Distinct roles for histone chaperones have also been suggested by recent structural and biochemical studies of their positions and interactions within the elongation complex^8,9,26,30–35^. However, how each histone chaperone contributes during transcription elongation remains a mystery.

While we do not understand why so many histone chaperones are required, there is growing evidence that several of them functionally interact during transcription, including FACT, Spt6, and Spn1/IWS1^35–39^. Consistent with this idea, depletion or mutation of FACT or Spt6 causes several common mutant phenotypes, suggesting that these two histone chaperones may be required for different steps of a common process^21,22,38,40–44^. Furthermore, Spt6 and Spn1/IWS1 physically interact^8,39,45,46^. Thus, the unwrapping and rewrapping of nucleosomes during transcription may require the coordinated activity of multiple essential histone chaperones.

Our studies focus on Spt6, an essential transcription elongation factor conserved throughout eukaryotes whose function as a histone chaperone is poorly understood^47^. While it is well-established that Spt6 binds directly to histones^39,48–50^, the region of Spt6 that interacts with histones and whether this interaction is important have been enduring questions. Spt6 is mostly comprised of well-structured domains but has a long, acidic, intrinsically disordered N-terminal domain (NTD)^8,32,51^. Two studies have suggested that the Spt6 NTD is required for histone binding in vitro^46,49^, but there is no direct evidence that this is the case, nor is there any evidence that an Spt6-histone interaction is important in vivo.

In this work, we uncover the function of the Spt6 NTD using a combination of genetic and biochemical approaches. First, we identify three acidic blocks within the Spt6 NTD that are together essential for viability. Second, by a combination of biochechemical approaches, including nuclear magnetic resonance (NMR), we show that specific regions of the NTD bind directly to all four histones, including the three acidic blocks identified in our genetic studies. Our NMR studies also reveal that Spn1/IWS1 and Elf1 regulate Spt6-histone binding. Third, we present evidence that the Spt6 NTD is both necessary and sufficient for histone binding in vivo, and that this activity is required for normal chromatin structure. Finally, we demonstrate that the requirement for the Spt6 NTD for viability and normal chromatin structure can be largely bypassed by a specific class of mutations in the histone chaperone FACT, illuminating a close functional relationship between the two. Taken together, our results define how Spt6 binds to histones and demonstrate that this binding performs a vital role in the cell. Furthermore, our results have led to a mechanistic model for how multiple essential histone chaperones coordinate the disassembly and reassembly of nucleosomes during transcription elongation.

## Results

### The Spt6 NTD is highly charged and essential for viability in *S. cerevisiae*

For our studies of Spt6-histone interactions, we focused on the N-terminal domain (NTD) of Spt6, which shares characteristics with many histone-binding domains: the Spt6 NTD is predicted to be intrinsically disordered, with the exception of the Spn1-binding region^26,45,46,51^, and it is highly charged, with an overall negative charge (Figure 1A-C)^7,52^. When we analyzed the distribution of charged residues in the Spt6 NTD, we found five acidic blocks interspersed with four basic blocks (Figure 1C). The human Spt6 NTD has a similar charge distribution, suggesting conservation of function^53^ (Figure S1A-C).

**Figure 1.**
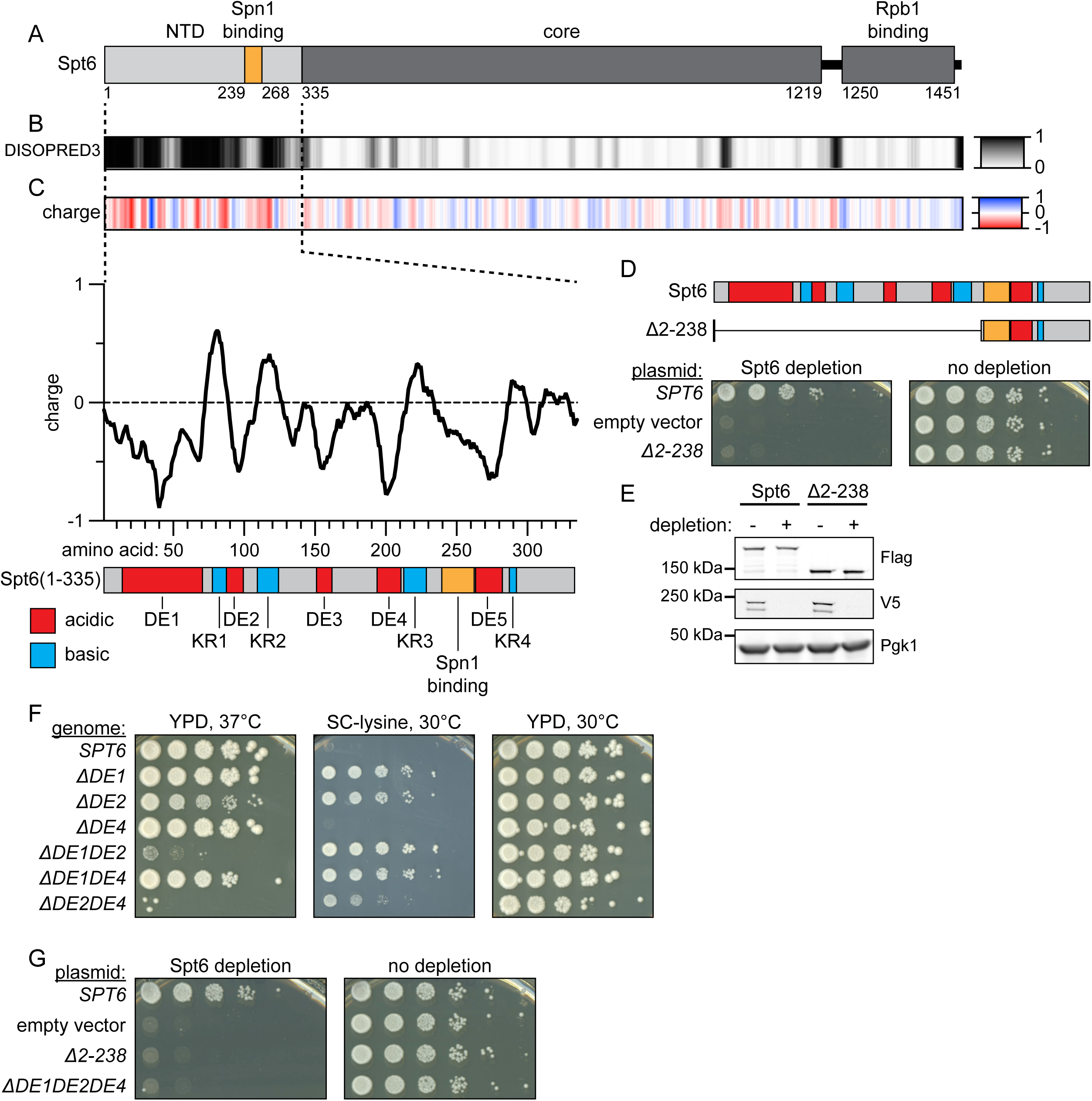
The Spt6 N-terminal domain is highly charged and essential for viability in *S. cerevisiae*. (A) Domain diagram of *S. cerevisiae* Spt6 depicting the N-terminal domain (NTD), core, and Rpb1 binding domains, with amino acid numbering underneath. The region of the NTD that binds Spn1 is highlighted in orange. (B) The disorder of Spt6 was predicted using DISOPRED3^76^. Results are shown as a heatmap with residues aligned to the domain diagram in (A). Higher values indicate a stronger predicted propensity for disorder. (C) Charge landscape of Spt6 depicted as a heatmap, showing the rolling average of amino acid side-chain charge in a fifteen-residue window. Below is a plot of the charge analysis for the Spt6 NTD (residues 1-335). Below the x-axis is a diagram of the Spt6 NTD showing the charged regions identified in this analysis. (D) The Spt6 NTD is essential for viability. At the top is a diagram depicting the NTD of the two versions of Spt6 used in this experiment. Below are spot tests showing whether the plasmid-encoded version of Spt6 can complement depletion of genome-encoded Spt6. (E) Western blot showing that Spt6Δ2-238 protein is present in undepleted and depleted conditions. (F) Three NTD acidic blocks are required for wild-type function. Spot tests were grown as indicated. Growth on SC-lysine shows expression of the *lys2-128δ* reporter^77^. (G) Three NTD acidic blocks are essential for viability. Spot tests for complementation of Spt6 depletion as in (D). See also Figure S1.

We first tested whether the Spt6 NTD is important for Spt6 function in vivo. We expressed either wild-type *SPT6* or a mutant lacking amino acids 2-238 (*spt6Δ2-238*) from plasmids and tested whether they could complement depletion of the wild-type Spt6 protein. We chose the 2-238 deletion as it removed seven of the nine NTD charged blocks (Figure 1D), and it was not predicted to disrupt the Spt6 Spn1-binding region (amino acids 239-268). Our results demonstrated that the strain expressing *spt6Δ2-238* was unable to grow upon depletion of wild-type, genome-encoded, Spt6 (Figure 1D-E). Therefore, the Spt6 NTD is required for viability in *S. cerevisiae*.

To test whether specific charged regions in the Spt6 NTD are required for Spt6 function, we constructed and analyzed two classes of mutations. First, we generated a series of nine *spt6* mutants in which all charged residues within each individual acidic or basic region were substituted with alanines (Figure S1D). We tested these nine mutants on plasmids for growth after depletion of wild-type Spt6 and found that all nine supported viability (Table S1). We then integrated each mutation into the genomic *SPT6* locus and screened the mutants for phenotypes previously associated with impaired Spt6 function. Three of the nine mutants, for regions DE1, DE2, and DE4, displayed mutant phenotypes (Table S1), while the other six did not. These results showed that three acidic regions of the Spt6 NTD are important for its function. However, additional analysis of the alanine-substitution mutations suggested that they caused neomorphic effects (Methods). Therefore, we constructed strains that contained genomic deletions of regions DE1, DE2, and DE4 (Figure S1E). These deletions caused phenotypes similar to the alanine-substitution mutations, although milder, particularly for DE4 (Figure 1F). All subsequent mutant analyses used the deletion alleles, as our results suggested that they most closely represent loss of function for each of the three NTD regions.

Given that *spt6Δ2-238* causes inviability, we tested for partial redundancy between Spt6 NTD regions DE1, DE2, and DE4. To do this, we constructed the three double deletion mutants and the triple deletion mutant (Figure S1E). For each double mutant, the strains were viable but had stronger mutant phenotypes than the single mutants, including strong temperature-sensitivity for two of the double mutants (Figure 1F). Furthermore, the triple deletion mutant, *spt6*Δ*DE1DE2DE4,* was inviable, similar to *spt6Δ2-238* (Figure 1G). Taken together, these results show that three specific acidic blocks in the Spt6 NTD contribute to a common essential function.

### The Spt6 NTD binds to histones in vitro

A key step in understanding how Spt6 functions as a histone chaperone is to understand whether the Spt6 NTD directly binds to histones. Using two related approaches, we tested whether the Spt6 NTD could form complexes with histones in vitro (Figure 2A). For these experiments, we purified two versions of the Spt6 NTD: the longer fragment (1-335) encompasses the complete NTD, including the Spn1-binding region, while the shorter fragment (1-238) includes the three acidic blocks identified in our genetic analysis as essential for Spt6 function.

**Figure 2.**
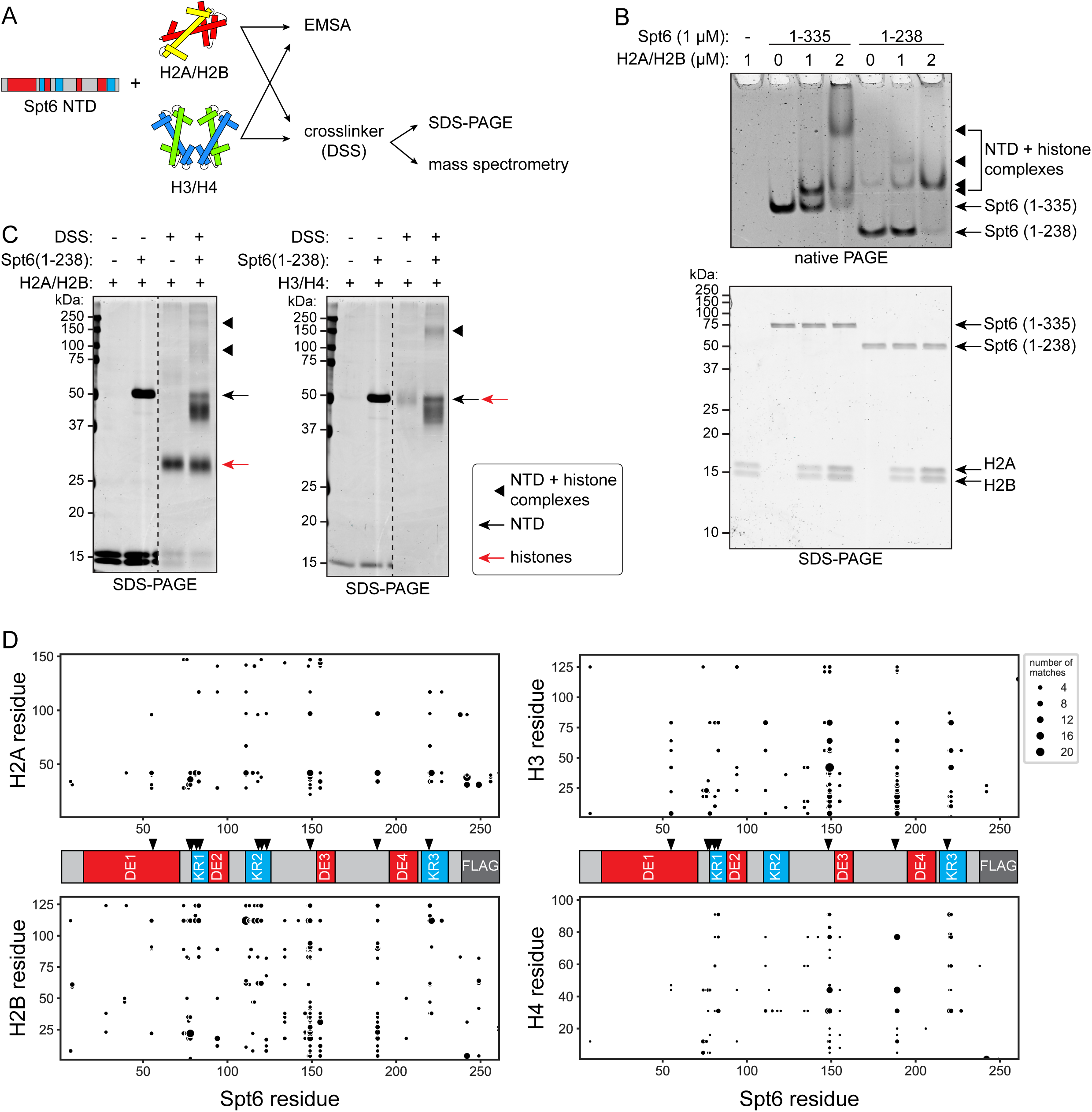
The Spt6 NTD binds to histones in vitro. (A) Schematic diagram of in vitro experiments. Purified recombinant Spt6 NTD constructs and histones were mixed and analyzed by electrophoretic mobility shift assay (EMSA) or by in vitro crosslinking followed by SDS-PAGE and mass spectrometry. (B) EMSA analysis of Spt6 NTD-H2A/H2B complexes. Purified Spt6 NTD fragments and histones H2A/H2B were mixed in the amounts indicated and separated by native or denaturing PAGE. Gels were stained with Coomassie to visualize proteins. The mobilities of different species are indicated. Western analysis suggested that the faint upper band in lane 5 of the native PAGE is an Spt6 (1-238) homodimer. (C) Crosslinking analysis of Spt6 NTD-histone complexes. Purified components were mixed as indicated prior to separation by denaturing PAGE. Gels were stained with Coomassie to visualize proteins. Intervening lanes have been removed for visualization purposes, as marked by the dashed line. The mobilities of different crosslinked complexes are indicated. Spt6 (1-238) alone does not form high molecular weight complexes (Figure S2B). (D) Mass spectrometry analysis of crosslinked Spt6 NTD-histone complexes. Crosslinked residues between the Spt6 construct and each histone are depicted in the scatter plots. The size of each dot indicates the number of times that each crosslink was observed. A schematic of the Spt6 NTD construct used is also shown, with acidic (DE, red) and basic (KR, blue) charge blocks indicated, as well as the Flag-tag used for protein purification. Black arrowheads indicate lysine residues that showed extensive crosslinks. See also Figure S2.

First, we tested for Spt6 NTD-histone binding by an electrophoretic mobility shift assay (EMSA) using native gel electrophoresis^50^. Under the conditions used, the positively-charged histones only migrate through the native gels when bound to the negatively-charged Spt6 NTD. Our results showed that H2A/H2B dimers formed stable complexes with both Spt6 (1-335) and Spt6 (1-238) (Figure 2B). Furthermore, we observed two distinct Spt6 (1-335)-H2A/H2B complexes, suggesting that a single Spt6 (1-335) molecule can bind to one or two H2A/H2B dimers. We did not observe a second higher molecular weight complex for the Spt6 (1-238)-H2A/H2B reactions, but the predicted Spt6 (1-238)-2(H2A/H2B) complex would not enter the gel under our electrophoresis conditions. Similarly, for H3/H4, we did not observe Spt6 NTD-H3/H4 complexes by EMSA, but we did observe that both the Spt6 (1-335) and Spt6 (1-238) bands disappeared with increasing amounts of H3/H4, suggesting that Spt6 NTD-H3/H4 complexes formed, but had an insufficient negative charge for gel migration under our electrophoresis conditions (Figure S2A).

Given our EMSA results, we took a second approach, chemical crosslinking, that would allow detection of Spt6 NTD-histone complexes regardless of their charge. We performed these experiments with only the Spt6 1-238 NTD fragment as it contained the three acidic regions identified by our genetic results and our EMSA results suggested that it was sufficient for histone binding. For this experiment, we again mixed the Spt6 NTD and either histones H2A/H2B or H3/H4, but this time we included disuccinimidyl suberate (DSS), a homobifunctional lysine-reactive crosslinker. The resulting complexes were then analyzed by both denaturing polyacrylamide gel electrophoresis and mass spectrometry (MS). When H2A/H2B or H3/H4 were incubated with DSS in the absence of the Spt6 NTD, we observed complexes with electrophoretic mobilities consistent with H2A/H2B dimers and (H3/H4)_2_ tetramers, respectively (Figure 2C). When Spt6 (1-238) was included, we observed higher molecular weight complexes, whose electrophoretic mobility indicated Spt6 (1-238)-H2A/H2B and Spt6 (1-238)-(H3/H4)_2_ complexes (Figure 2C; S2B). MS analysis revealed extensive crosslinks between Spt6 (1-238) and all four histones (Figure 2D; S2C-D; Table S2). Notably, Spt6 residues K55, K77, K78, K81, K83, K149, K189, and K220 formed crosslinks with both H2A/H2B and H3/H4, while Spt6 residues K118, K120, and K123 only formed substantial crosslinks with H2A/H2B. These results, taken together with our EMSA results, demonstrate that similar regions of the Spt6 NTD can bind to histones H2A/H2B and H3/H4 in vitro and that Spt6 amino acids 1-238 are sufficient for these interactions.

### NMR reveals that specific regions of the Spt6 NTD bind directly to histones

To better resolve which regions of the Spt6 NTD bind directly to histones, we used nuclear magnetic resonance (NMR) spectroscopy. In these experiments, we tested both Spt6 (1-238) and Spt6 (1-335). When we probed the binding of H2A/H2B or H3/H4 with each Spt6 NTD fragment, we observed a substantial loss of peak intensity for multiple residues in Spt6 (example shown in Figure 3A-C; Methods), indicating specific interactions between the Spt6 NTD and both sets of histones. The NMR spectra revealed similar binding patterns for H2A/H2B (Figure 3D) and H3/H4 (Figure 3E), consistent with our crosslinking results. In addition, while Spt6 (1-238) showed generally stronger binding than did Spt6 (1-335), the 1-335 fragment included an additional binding region that is not present in Spt6 (1-238), an acidic region known to bind Spn1 (amino acids 239-268)^45,46^. This could explain previous reports that Spt6 residues 239-298 are important for histone binding in vitro^49^. Strikingly, the three acidic blocks identified by our genetic results as important for Spt6 NTD function were regions of direct interaction between Spt6 and histones.

**Figure 3.**
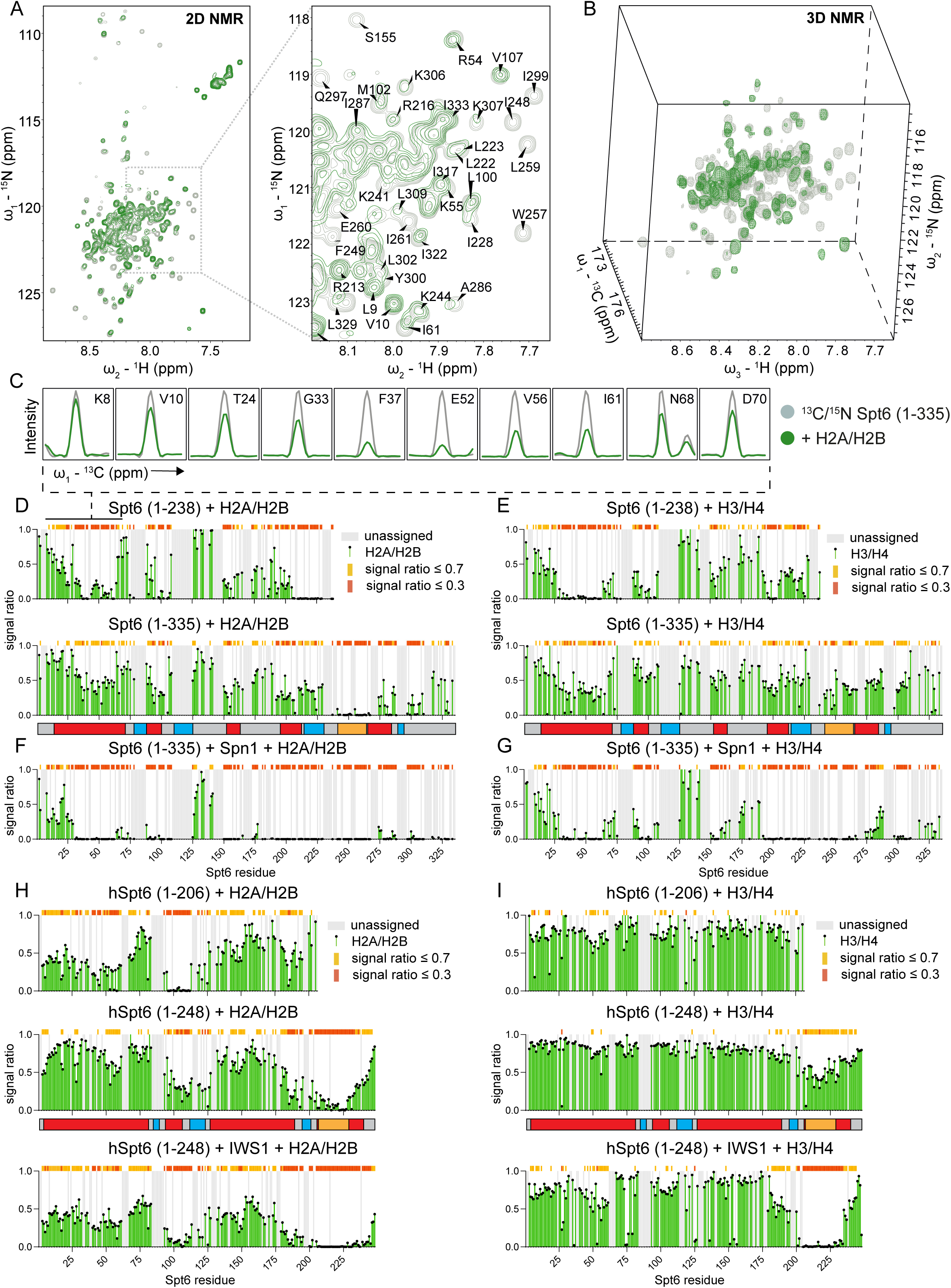
NMR reveals that specific regions of the Spt6 NTD bind directly to histones. (A-C) Overlaid 2D ^15^N/^1^H HSQC spectra obtained for free (gray) or H2A/HB-bound (green) ^15^N labeled Spt6 (1-335). Signals in certain regions within the 2D NMR spectra of intrinsically disordered regions are highly overlapped, resulting in information loss. To address this, we followed the interactions in less overlapped 3D spectra, as shown in panel (B). Panel (C) provides an example of this analysis, with cross-sections from the free (gray) and H2A/H2B-bound 3D HNCO spectra of ^13^C/^/15^N labeled Spt6 (1-335, green). Binding is manifested by a reduction in signal intensity for specific Spt6 residues, with near-zero values suggesting strong interactions. (D and E) The plots of the NMR signal intensity ratios obtained for Spt6 NTD constructs upon H2A/H2B (D) and H3/H4 (E) binding. Residues with signal ratios below 0.3 or 0.7, indicating strong or medium binding, are highlighted as indicated. Residues that could not be assigned are shown in gray. A schematic of the charged regions of the Spt6 NTD is aligned with the x-axes of the plots, including acidic (red), basic (blue) and Spn1-binding (orange) regions. (F and G) As in (D and E), but with the addition of Spn1. (H and I) As above, but for labeled human Spt6 NTD with histones H2A/H2B (H) or H3/H4 (I) and IWS1, the human homolog of *S. cerevisiae* Spn1. A schematic of the charged regions of the human Spt6 NTD is aligned with the x-axes of the plots, including acidic (red), basic (blue) and IWS1-binding (orange) regions. See also Figures S3 and S4.

To determine the regions of histones that are bound by Spt6, we titrated ^15^N labeled histones H2A/H2B or H3/H4 with Spt6 peptides and followed the binding by their NMR spectra. The Spt6 peptides corresponded to distinct regions of the NTD and represented most of the regions for which we observed strong binding to histones in our Spt6 NTD experiments (Figure S3A). Using peptides instead of the entire Spt6 NTD allowed us to individually map the histone surfaces bound by different Spt6 histone binding regions. From these experiments we obtained interpretable spectra for histones H2A/H2B, but not H3/H4, due to the heterogeneity linked to the H3/H4 dimer-tetramer equilibrium (data not shown). For H2A/H2B, the results indicated that the Spt6 peptides bound to the same H2A/H2B surface that binds to DNA (Figure S3B-F). These results suggested that multiple regions of the Spt6 NTD bind to a common surface on the H2A/H2B dimer.

Given the conservation of amino acid sequence and charge distribution between the yeast and human Spt6 NTDs, we also used NMR to test whether the human Spt6 NTD binds to histones. We used human Spt6 (1-206) and human Spt6 (1-248), corresponding to yeast Spt6 (1-238) and (1-279), respectively. The NMR spectra for the human Spt6 NTDs showed substantial interactions with histones H2A/H2B (Figure 3H), although only weak interactions with histones H3/H4 (Figure 3I). This result suggests that histone binding is a conserved function of the Spt6 NTD.

### Spn1 and Elf1 contribute to Spt6-histone binding

Recent studies have shown that Spt6 can form a complex with Spn1 and either H2A/H2B or H3/H4^39^. Therefore, we tested whether Spn1 alters Spt6-histone interactions as detected by NMR. First, we probed Spt6 (1-335) binding to Spn1 alone and observed the expected strong binding over the previously established Spn1-binding region (Figure S4A, row 1). When we incubated Spt6 (1-335) with Spn1 plus either histones H2A/H2B or H3/H4, we observed that Spn1 substantially increases binding between Spt6 (1-335) and histones, although we cannot rule out that some of the changes in peak intensity are due to Spt6-Spn1 interactions (Figure 3F-G; S4A, rows 2, 3, 8, 9). We also observed a similar effect on human Spt6 (1-248) with both H2A/H2B and H3/H4 when the human homolog of Spn1, IWS1, was included (Figure 3H-I). These results suggest a conserved regulatory capacity for Spn1/IWS1 to stimulate Spt6 NTD-histone binding.

To more extensively characterize possible Spt6-Spn1-histone complexes, we also investigated binding to Spn1 by both histones and Spt6. For full-length Spn1, we detected signals only from the dynamic segments that could be assigned to the N-terminal region of Spn1 (Figure S4B). As expected, those results showed strong binding to histones H3/H4 and weak binding to H2A/H2B^39^ (Figure S4B, rows 5, 8). We also examined binding to a version of Spn1 lacking its N-terminal domain, which allowed analysis of the Spn1 conserved core and C-terminal region (Figure S4C). Again, we observed preferential binding to histones H3/H4 (Figure S4C, rows 2, 6), indicating a more extensive Spn1-H3/H4 interaction than previously reported^39^. The binding of H3/H4 to Spn1 was diminished by the addition of the full Spt6 NTD (1-335) but remained unaffected upon addition of Spt6 (238-269), which interacts predominantly with the structured Spn1 core^45,46^ (Figure S4C, rows 6, 7, 10, 12). These observations are consistent with previous results showing that in a complex of Spn1-Spt6-H3/H4 histones preferentially bind to Spt6 and not to Spn1^39^.

We also investigated another elongation factor, Elf1^27^, as it directly interacts with Spn1^8,26,54^. Elf1 increased Spt6-histone interactions, particularly for histones H2A/H2B (Figure S4A, rows 2, 6). For H3/H4, the Elf1-stimulated increase appeared primarily over the DE1 region of the Spt6 NTD (Figure S4A, rows 8, 10). Analysis of binding to Elf1 itself showed that it could bind histones H2A/H2B and H3/H4, although this interaction was greatly diminished by the addition of the Spt6 NTD, suggesting that a main role of Elf1 is to stimulate the binding of histones to Spt6 (Figure S4D, rows 1, 3, 9, 11).

Our NMR results and previous genetic results^27^ suggested that both Spn1 and Elf1 are important for Spt6 function in vivo. To test this for *spt6* NTD mutations, we constructed a set of *spt6 spn1* and *spt6 elf1* double mutants to assay whether *spn1* or *elf1* mutations would exacerbate any Spt6 NTD mutant phenotypes. For the essential *SPN1* gene we used a missense mutation, *spn1-F267E*^46^, and for *ELF1* we used an *elf1* deletion. Our results showed that both the *spt6 spn1* and *spt6 elf1* double mutants had considerably more severe phenotypes than any of the single mutants, including lethality for the *spt6 elf1* double mutants (Table S3). The NMR results, taken together with these genetic interactions, suggest that specific regions of the Spt6 NTD, Spn1, and Elf1 contribute to a common essential function in vivo: the binding of histones to Spt6.

### The Spt6 NTD is necessary and sufficient for Spt6-histone interactions in vivo

To directly test whether the Spt6 NTD binds to histones in vivo, we performed coimmunoprecipitation (coIP) experiments comparing wild-type Spt6 to the Spt6Δ2-238 mutant. To do this, we expressed either full-length Spt6 or Spt6Δ2-238 fused to a Flag epitope tag and then performed coIPs in strains also expressing wild-type Spt6. We observed coIP of full length Spt6 and all four core histones (Figure 4A). In contrast, in the Spt6Δ2-238 mutant, coIP was undetectable for all four histones, demonstrating that the Spt6 NTD is necessary for Spt6-histone interactions in vivo (Figure 4A).

**Figure 4.**
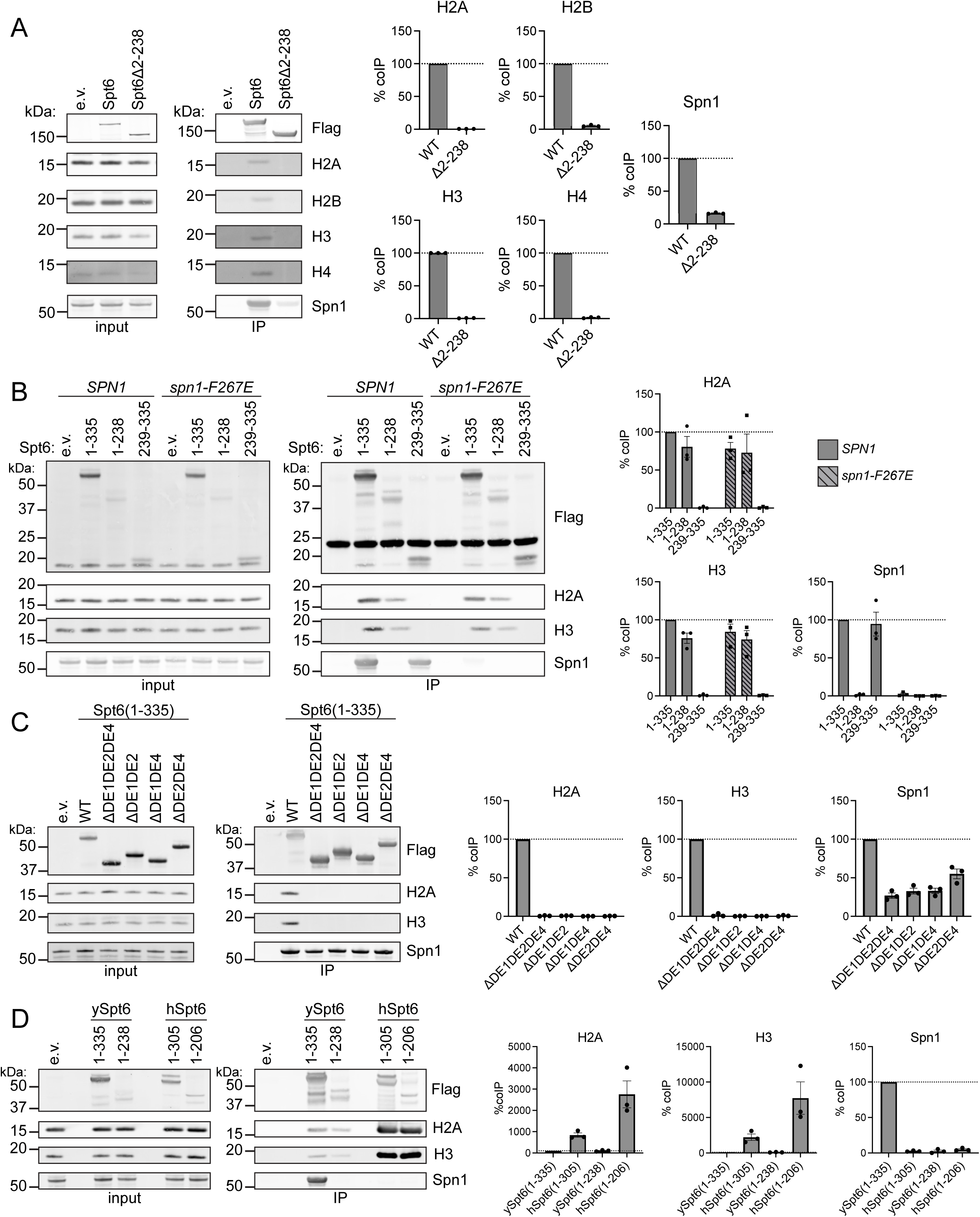
The Spt6 NTD is necessary and sufficient for Spt6-histone interactions in vivo. (A-D) Coimmunoprecipitation of indicated proteins with Flag-tagged Spt6 constructs. Quantification of three biological replicates is shown to the right of each panel, with data represented as the mean ± SEM. (A) Spt6 residues 2-238 are required for histone binding. WT: Spt6; Δ2-238: Spt6Δ2-238. (B) Spt6 residues 2-238 are sufficient for histone binding in wild-type (*SPN1*) or mutant (*spn1-F267E*) cells. (C) Three acidic blocks in the Spt6 NTD are required for histone binding. (D) Histone binding is a conserved function of *S. cerevisiae* (ySpt6) and human (hSpt6) Spt6 NTDs.

To test whether the Spt6 NTD is sufficient for binding to histones in vivo, we expressed three different Spt6 NTD fragments in cells (1-335, 1-238, and 239-335) and measured their coIP with histones H2A and H3. Our results showed that the histones coIP with Spt6 (1-335) and Spt6 (1-238), but not with Spt6 (239-335), which contains the Spn1 binding region (Figure 4B). Thus, Spt6 (1-238), which contains three regions that bind histones in vitro, is both necessary and sufficient for binding histones in vivo.

Previous results^8,39^, along with our NMR results, suggested that Spn1 contributes to the Spt6-histone coIP that we observed. To test this possibility, we repeated the Spt6 NTD-histone coIP experiments in a *spn1-F267E* mutant, in which the Spt6-Spn1 interaction is undetectable^46^. Our results showed that in the *spn1-F267E* mutant, H2A and H3 were able to coIP with Spt6 (1-335) and Spt6 (1-238) at levels similar to those in the wild-type background (Figure 4B). As a control, we showed that Spn1 was able to coIP with both Spt6 (1-335) and Spt6 (239-335) in a wild-type background but not in the *spn1-F267E* mutant (Figure 4B). We conclude that the Spt6 NTD is sufficient for interaction with histones in vivo and that this interaction is Spn1-independent, although Spn1 may perform roles in vivo not captured by our coIP conditions.

We next set out to determine whether the three acidic regions that we had identified by genetic analysis and NMR were required for histone binding. To do this, we constructed the three possible double deletion mutants and the triple deletion mutant in the Spt6 (1-335) expression construct and analyzed Spt6 NTD-histone interactions by coIP. Our results demonstrated that there was no detectable Spt6 NTD-histone coIP for any of these NTD mutants (Figure 4C), showing that these three regions are necessary for Spt6 NTD-histone interactions. The lack of Spt6 NTD-histone interactions in any of the double deletion mutants shows that no individual acidic region is sufficient for Spt6-histone coIP. We also found that the level of Spn1 coIP was reduced in the mutants, with the triple deletion allele having the greatest effect (Figure 4C). This result was consistent with the reduced level of Spt6-Spn1 coIP observed in the *spt6*Δ*2-238* mutant (Figure 4A) and with a previous study that showed that eight Spt6 phospho-serines distributed throughout the Spt6 NTD contribute to the Spt6-Spn1 interaction^55^. In conclusion, our results establish that three in vitro histone binding interfaces of the Spt6 NTD, as identified by NMR and mutant analysis, are required for Spt6-histone interactions in vivo.

Finally, we tested whether the conserved human Spt6 NTD could bind to histones in vivo in *S. cerevisiae.* To do this, we expressed two human Spt6 NTDs (1-305) and (1-206) in yeast, corresponding to yeast Spt6 NTDs (1-335) and (1-238), respectively, and assayed their interactions with histones by coIP. Surprisingly, our results showed that the human Spt6 constructs resulted in substantially greater histone coIP than their yeast counterparts (Figure 4D). Neither human Spt6 construct bound to yeast Spn1 (Figure 4D). The increased level of histone binding for the human Spt6 NTD compared to the yeast Spt6 NTD may be due to the greater negative charge of the human NTD (Figure S1A-B). These results demonstrate that histone binding is a conserved function of the Spt6 NTD.

### The Spt6 NTD is not required for the association of Spt6 with chromatin

Previous results showed that the association of Spt6 with chromatin requires the Spt6 tSH2 domain, which interacts with RNAPII^54,56^. However, it has never been tested whether the association of Spt6 with chromatin also requires Spt6-histone interactions. To test this possibility, we performed chromatin immunoprecipitation-sequencing (ChIP-seq), comparing full-length Spt6 to Spt6Δ2-238 (Figure 5A; S5F). Our results showed that, prior to depletion, Spt6Δ2-238-Flag did not ChIP at levels comparable to full-length Spt6-Flag (Figure S5E). However, after depletion of genome-encoded Spt6-V5, Spt6Δ2-238-Flag bound to chromatin at levels similar to full-length Spt6-Flag (Figure 5B; S5B,C,E). The global level of RNAPII occupancy was also largely unaffected in the mutant (Figure 5C), and, as expected, the level of Spt6 occupancy correlated with the level of RNAPII occupancy (Figure 5D). Our results demonstrate that the NTD is not required for normal levels of Spt6 association with chromatin.

**Figure 5.**
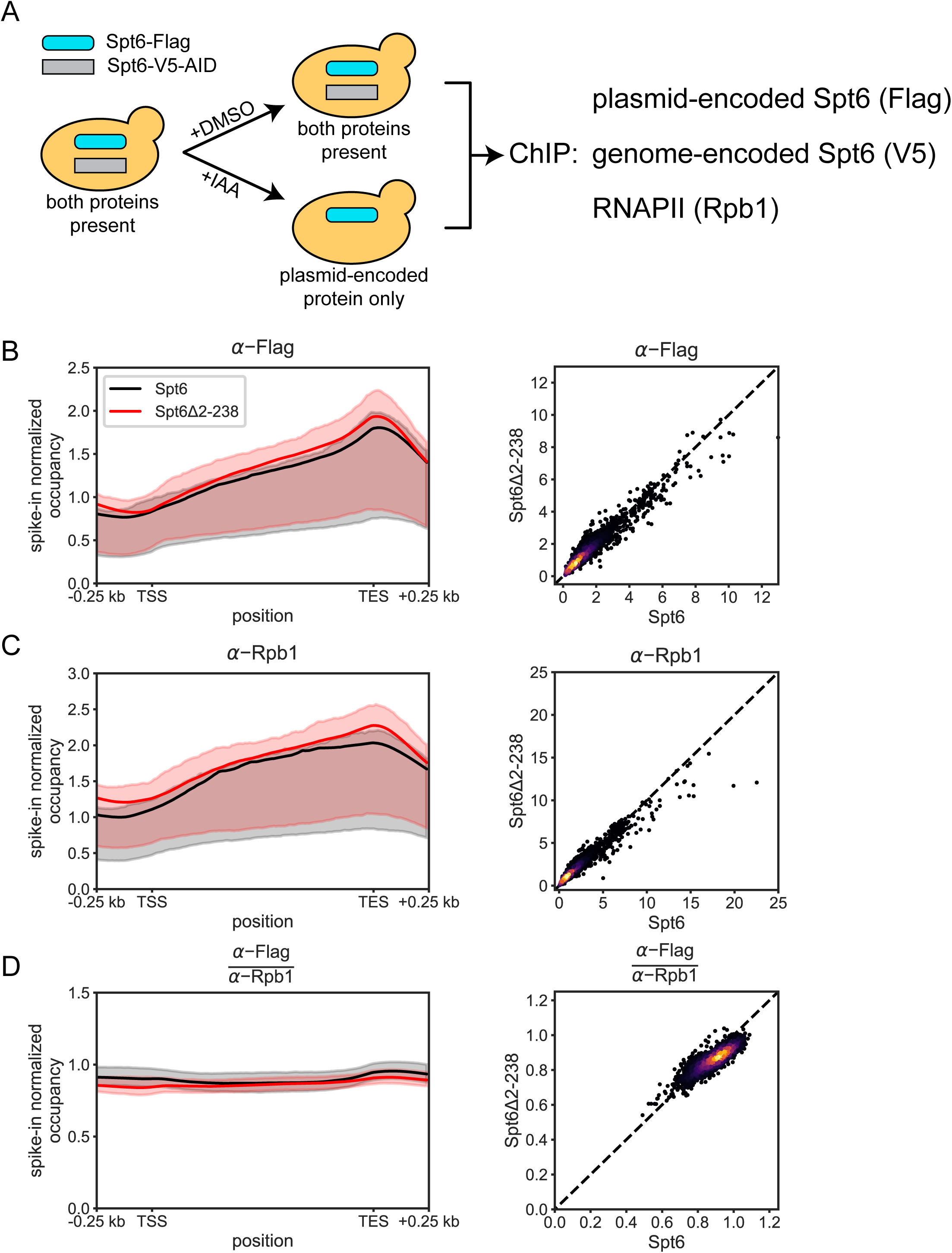
The Spt6 NTD is not required for the association of Spt6 with chromatin. (A) A schematic of the conditions analyzed by ChIP-seq. (B) Left: a metagene plot of Flag-tagged Spt6 (black) and Spt6Δ2-238 (red) spike-in normalized ChIP-seq occupancy for 3,087 non-overlapping protein-coding genes scaled to the same length. The line represents the average occupancy at each position calculated in bins of 10 bp. The shadings indicate the inter-quartile ranges, which are shown to represent the diversity of genes plotted. Right: a scatterplot of per-gene occupancy values for Spt6Δ2-238 cells vs. Spt6 cells. Each dot represents the average ChIP occupancy of a gene represented in the metagene. The dashed line plots *y = x*. Dots are colored using a gaussian kernel density estimate to depict the density of the data. Shown is a representative of three biological replicates (replicate 3; see Figure S5A-C). (C) As in (B), but for Rpb1 (8WG16) ChIP-seq occupancy. (D) As in (B), but for Spt6 ChIP-seq occupancy normalized by Rpb1 occupancy. See also Figure S5.

### The Spt6 NTD is required for normal chromatin structure in vivo

Previous tests of the requirement of Spt6 for normal chromatin structure examined the consequences of Spt6 depletion, which is known to alter the composition of the transcription elongation complex^37,39,57–60^. Studying the Spt6Δ2-238 mutant provides the opportunity to test whether the Spt6 NTD is required for normal chromatin structure while minimizing or eliminating the major perturbations caused by depletion of the entire Spt6 protein. In this way, we can determine the extent to which normal chromatin structure depends specifically on the histone-binding activity of Spt6. Therefore, we used our Spt6 depletion system (Figure 5A) and performed micrococcal nuclease sequencing (MNase-seq) of cells expressing either wild-type Spt6, Spt6Δ2-238, or no Spt6. The wild-type Spt6 condition displayed a normal nucleosome pattern, with well-positioned +1 nucleosomes, followed by regularly phased nucleosome arrays within gene bodies (Figure 6A-B; S6C). When we depleted Spt6, we observed a genome-wide decrease in nucleosome occupancy as well as a progressive 3’ shift in nucleosome position within gene bodies (Figure 6A-B), similar to previous Spt6-depletion studies^19,44^. The *spt6Δ2-238* mutant also showed genome-wide alterations in both nucleosome occupancy and positioning, albeit with a less severe decrease in occupancy and 3’ shift of nucleosome positioning than observed in Spt6 depletion conditions (Figure 6A-B). These results demonstrate that the Spt6 NTD is required for normal nucleosome organization throughout the genome.

**Figure 6.**
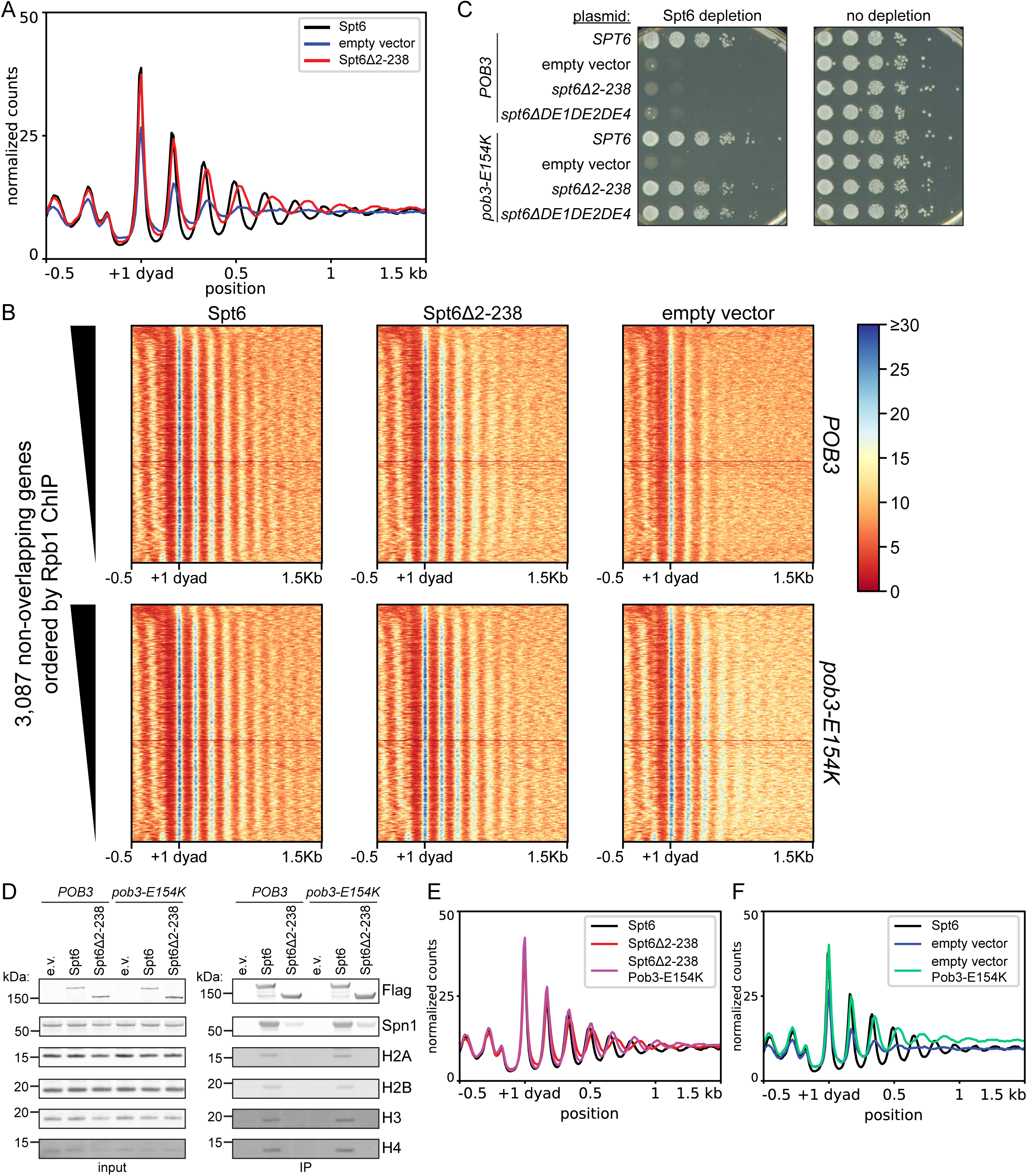
The Spt6 NTD and FACT cooperate to control chromatin structure. (A) The Spt6 NTD is required for normal chromatin structure. Metagene plot of spike-in normalized nucleosome dyad occupancy in non-overlapping 20 bp bins in cells expressing wild-type Spt6 (black), Spt6Δ2-238 (red), or no Spt6 (empty vector; blue). The lines represent the mean occupancy across 3,087 non-overlapping protein-coding genes. Shown is a representative of three biological replicates (replicate 3; see Figure S6B). (B) Heatmaps showing spike-in normalized nucleosome occupancy and positioning at 3,087 non-overlapping protein-coding genes ordered by decreasing Rpb1 ChIP-seq occupancy. Heatmaps are arrayed in columns and rows according to *SPT6* and *POB3* alleles, respectively. (C) A change in the FACT subunit Pob3 bypasses the requirement for the Spt6 NTD. Spot tests for complementation of Spt6 depletion as in Figure 1G. (D) The *pob3-E154K* mutation does not restore Spt6 binding to histones. CoIP of the indicated proteins with Flag-tagged Spt6 or Spt6Δ2-238 in wild-type (*POB3*) or mutant (*pob3-E154K*) cells. Images from the same replicate are shown in Figure 4A. Quantification of biological replicates is shown in Figure S6A. (E) Spt6Δ2-238-associated chromatin structure defects are suppressed by *pob3-E154K*. Metagene plot is as in (A). (F) Chromatin structure defects associated with total loss of Spt6 are partially improved in *pob3-E154K* cells. Metagene plot is as in (A). See also Figure S6.

### Mutations that alter FACT bypass the requirement for the Spt6 NTD

Previous studies have suggested that multiple histone chaperones act in a coordinated fashion^37,42,61^. To identify factors that functionally interact with the Spt6 NTD, we selected for suppressors of the temperature-sensitive phenotype of the *spt6*Δ*DE2DE4* mutant. We isolated six independent, recessive mutants that were able to grow at the non-permissive temperature (37°C) and identified the causative mutations by whole-genome sequencing and genetic analysis (Methods). Our results showed that three of the mutations were in *SPT16* and three of the mutations were in *POB3*, the genes that encode the two subunits of the histone chaperone complex FACT (Table S4). The suppressor mutations were located near previously isolated *spt16* and *pob3* mutations that bypass the requirement for Spn1^36,37^. Therefore, we tested a previously characterized mutation, *pob3-E154K*, and found that it strongly suppressed *spt6*Δ*DE2DE4*. Strikingly, *pob3-E154K* also strongly suppressed the inviability caused by an NTD deletion, *spt6Δ2-238* (Figure 6C; Table S3). Thus, changes in the histone chaperone FACT bypass the essential requirement for the Spt6 NTD. This suppression is specific for the Spt6 NTD, as *pob3-E154K* does not suppress the complete loss of Spt6 (Figure 6C).

The previous mutant selections that discovered *pob3-E154K* also identified strong suppressor mutations in several other genes^36,37^. To determine the extent of similarity between bypass of the Spt6 NTD and bypass of Spn1, we tested two other classes of strong Spn1 bypass suppressors, deletion of *SET2* (which encodes an H3K36 methyltransferase) and deletion of *CHD1* (which encodes an ATP-dependent chromatin remodeler), and found that these deletions are extremely weak suppressors of *spt6Δ2-238* (Table S3). We also tested overexpression of *SPN1*, previously shown to suppress *spt6-YW,* a mutation that abolishes the Spt6-Spn1 interaction^37^. Our results showed that *SPN1* overexpression does not suppress *spt6Δ2-238* (Table S3). Together, these results suggest that strong bypass of the Spt6 NTD is limited to changes in FACT.

One possible mechanism for suppression of *spt6Δ2-238* by *pob3-E154K* is that the suppressor mutation restores Spt6-histone interactions. To test this possibility, we performed Spt6-histone coIP experiments in a *pob3-E154K* background. Our results showed that there was still no detectable coIP between Spt6Δ2-238 and any of the four core histones in the *pob3-E154K* background (Figure 6D; S6A). This result demonstrates that bypass of the Spt6 NTD by *pob3-E154K* includes bypass of the requirement for Spt6-histone interactions.

To test whether suppression of *spt6Δ2-238* by *pob3-E154K* occurs at the level of chromatin structure, we performed MNase-seq in a *pob3-E154K* background. Our results showed that *pob3-E154K* strongly suppressed the chromatin defects caused by the *spt6Δ2-238* mutation (Figure 6B,E; S6D). Interestingly, the chromatin defects caused by depletion of Spt6 were also weakly suppressed by *pob3-E154K* (Figure 6B,F). The ability of FACT mutants to compensate for defects in Spt6 suggests that these two histone chaperones cooperate to regulate chromatin structure genome wide.

## Discussion

Our results have provided substantial advances in addressing the longstanding question of how histone chaperones are utilized during transcription elongation through a nucleosome. First, we have shown that the Spt6 NTD is both necessary and sufficient for Spt6-histone interactions in vivo, and that these interactions depend upon three acidic blocks within the NTD. We also found that Spt6-histone interactions are critical in vivo as an Spt6 mutant missing these three acidic blocks does not support viability and loss of the Spt6 NTD changes chromatin structure across the genome. Agreeing with our in vivo results, our in vitro results showed that the Spt6 NTD binds directly to histone H2A/H2B dimers and H3/H4 tetramers. Furthermore, our NMR studies revealed that this binding occurs over specific regions of the Spt6 NTD that largely align with the three acidic blocks identified in vivo. The convergence of our genetic and biochemical results strongly suggests that the in vivo function of the three acidic blocks in the Spt6 NTD is to directly bind histones, a requirement for histone chaperone function. Finally, we have obtained enlightening evidence for a close functional relationship between Spt6 and FACT by the isolation of FACT mutants that bypass the inviability and chromatin defects caused by the loss of the Spt6 NTD. Taken together, our results suggest a dynamic mechanism by which Spt6 functions coordinately with FACT and other histone chaperones during transcription elongation through a nucleosome.

Three aspects of Spt6-histone binding have provided clues to how Spt6 functions as a histone chaperone: first, multiple regions of the Spt6 NTD bind to histones; second, the same regions of the Spt6 NTD bind to both histones H2A/H2B and H3/H4; and third, these regions of the Spt6 NTD likely bind to H2A/H2B on the DNA-binding surface of the histone dimer. Together, these results suggest that the Spt6 NTD adopts multiple histone binding configurations during transcription elongation by engaging histone surfaces normally occupied by nucleosomal DNA.

Here we present a model for how the histone binding activities of Spt6 and FACT coordinate a step-by-step process during transcription through a nucleosome. This model integrates the results presented in this paper, AlphaFold predictions^62^ of the binding of the Spt6 NTD to histones (Figure S7), genetic and biochemical studies of FACT^10,12,42,61,63–69^, and structural studies of FACT and the RNAPII elongation complex^3,8,9,11,30,70^. This model also accounts for the long-standing observations that Spt6 is tightly associated with RNAPII during elongation, while, in contrast, FACT is only recruited to the elongation complex once a nucleosome is partially unwrapped. Our model, illustrated in Figure 7, proposes that, as elongating RNAPII begins to unwind the nucleosome, the Spt6 NTD initiates the chaperoning process by mimicking DNA and binding to the proximal H2A/H2B dimer on its newly exposed surface (Figure 7, step 2; S7A). Then, as transcription proceeds and further destabilizes the nucleosome, FACT is recruited^9,11,64,65^, and the Spt6 NTD is displaced from the DNA-binding surface of the H2A/H2B dimer while remaining engaged with the face of H2B (Figure 7, step 3). Transcription then fully unwraps the nucleosome, resulting in the complete separation of the histones from DNA. FACT remains bound to the histones and the Spt6 NTD occupies the vacant DNA-binding surface of the H3/H4 tetramer (Figure 7, step 4; S7B), thus retaining the histones at the site of transcription and preventing their loss. This step is followed by a series of histone binding and release steps by Spt6 and FACT, coordinated to allow the histones to reassociate with DNA in an ordered fashion behind RNAPII (Figure 7, step 5). These actions culminate in the reassembly of the nucleosome upstream of the elongation complex (Figure 7, step 6).

**Figure 7.**
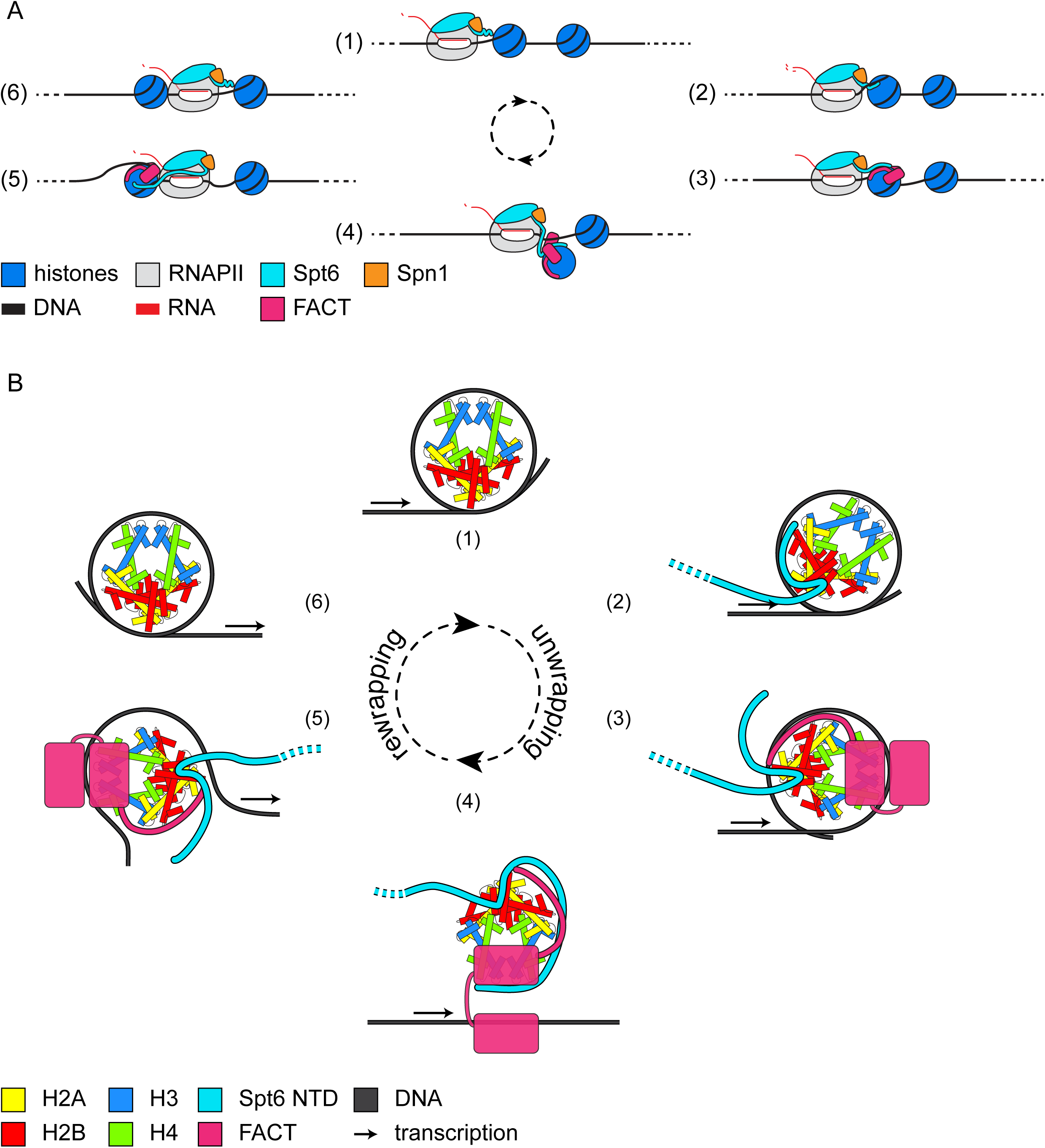
A model for co-transcriptional histone transfer. (A) A depiction of RNAPII transcribing through a nucleosome with Spt6, Spn1, and FACT. (1) Elongating RNAPII begins to unwind the nucleosome, breaking contacts between DNA and the proximal H2A/H2B dimer. (2) The Spt6 NTD binds to the proximal H2A/H2B dimer in two places, DE1 binds along the DNA-binding surface and DE2 binds the face of H2B (see Figure S7A). (3) Transcription further destabilizes the nucleosome, recruiting FACT^11,64,65^. The Spt16 CTD displaces Spt6 NTD DE1, but not DE2 from the proximal H2A/H2B DNA-binding surface. DE2 binding may maintain the association between the Spt6 NTD and the nucleosome to promote subsequent steps of the process. (4) DE1 of the Spt6 NTD replaces DNA on the H3/H4 tetramer, allowing the histones to fully dissociate from DNA while remaining tethered by both FACT and Spt6. A similar intermediate has recently been observed between histones, FACT, and the acidic IDR of Mcm2 during replication^71^. (5) After RNAPII passage, the nucleosome begins to be reassembled upstream of the elongation complex. DE1 of the Spt6 NTD is again replaced by DNA on the surface of the H3/H4 tetramer. FACT remains bound to DNA, the H3/H4 tetramer, and the proximal H2A/H2B dimer. DE2 may also remain bound to the proximal H2A/H2B dimer or may disengage so that the Spt6 NTD is available for transcription of the downstream nucleosome. (6) Transcription through the nucleosome is complete as the elongation complex moves to the next nucleosome. The steps described are based on structures from^8^ as follows: (2) EC42, PDB 7XSE; (3) EC49, PDB 7XSX; (5) EC58, PDB 7XTD; (6) EC115, PDB 7XSZ. Step (4) is proposed based on the structure of FACT during replication reported in^71^, see also Figure S7C. (B) Depiction of a more detailed view of the histone binding domains of Spt6 and FACT interacting with histones during nucleosomal transcription. The steps correspond to those depicted in (A). Placement of the Spt6 histone binding domains is based on our results and Alphafold 3 predictions^62^ (see Figure S7A-B). The position of RNAPII is indicated by the arrow. Recent work has suggested that hexasomes may result from loss of one H2A/H2B dimer during transcription elongation^8,9,65,78^. Our model does not currently distinguish between the formation of a hexasome or a nucleosome in the wake of transcription. See also Figure S7.

We propose additional facets of our model based on the bypass of the essential requirement for the Spt6 NTD by specific changes in FACT. The amino acid changes in the FACT mutants map to positions in Spt16 and Pob3 that likely reduce the level of association of FACT with nucleosomal DNA or result in a more ‘open’ conformation of FACT (Figure S7C)^8,11,71^. In support of this, our previous studies showed that *pob3-E154K* reduces Pob3-H3 interactions and the level of association of FACT with chromatin^37^. Relevant to our model, *pob3-E154K* and similar mutations were initially identified as bypass suppressors of the loss of the essential histone chaperone Spn1^36,37^. Spn1 physically interacts with Spt6 and is positioned to interact with histones as the nucleosome unwraps^8^. These results, along with our NMR results, previous characterization of Spn1-histone interactions^39,72^, and structural studies of Spn1 in the elongation complex^8^, suggest that Spn1 contributes to histone chaperone activity during the steps diagrammed in Figure 7. We propose that a balance of histone chaperone activities is required between the activities of Spt6 plus Spn1 and the activity of FACT. By this model, when either the Spt6 NTD or Spn1 are lost and their combined chaperone activity is reduced, the level of FACT must also be reduced to maintain a proper balance of activity. We note that there is strong genetic evidence that the level of histone chaperones is crucial, as either elevated or decreased levels of Spt6 and Spt16 cause mutant phenotypes^13,40,73,74^. Our model predicts that *pob3-E154K* would be unable to bypass loss of both the Spt6 NTD and Spn1, as that would further disrupt the balance of histone chaperone activities. Indeed, *pob3-E154K* cannot suppress a combination of *spt6*Δ*2-238* and *spn1*Δ, while it can suppress each one individually (Table S3). Thus, there is a tight functional connection between the activities of Spt6, Spn1, and FACT.

Our model does not yet account for other members of the transcription elongation complex that are required for transcription through a nucleosome. A previous study suggested that Elf1 and Spt4/Spt5 (also known as DSIF), cooperate to help RNAPII elongate into a nucleosome^26^. A positive role for Elf1 is supported by our NMR results and there is substantial evidence that Spt5 functions as a histone chaperone during elongation through nucleosomes^26,58,59^. Based on genetic evidence, these two activities may well functionally interact with Spt6, Spn1, and FACT^27,40,74^. Finally, there is evidence that when either Spt6 or FACT are depleted, additional histone chaperones (Asf1, Hir, and Rtt106) are recruited to chromatin^42,61^. Conceivably, such recruitments occur in some of the mutant conditions studied here.

In addition to the role that we establish in histone binding, the Spt6 NTD is an IDR that is part of a network of factors that co-localize within the nucleus^35^. It has been shown that the charge distribution of the Spt6 IDR is both required and sufficient for selective partitioning, possibly into nuclear condensates^53,75^. Given the overlap between the requirements for histone binding and selective partitioning by the Spt6 NTD, it would be interesting to determine whether histone binding is required for or drives selective partitioning of Spt6 and other proteins in the nucleus.

In conclusion, our experiments have established histone binding as a specific molecular function of the intrinsically disordered Spt6 NTD. Our findings clearly implicate collaboration between the histone chaperones Spt6, Spn1, and FACT as crucial for this process. How these factors coordinate binding at the scale of the nucleosome and how binding at this scale affects global chromatin structure remains an open question. Further structural, biochemical, and genetic characterization of these factors is needed to fully understand this essential, intricate process.

## SUPPLEMENTAL INFORMATION

provided for review as a PDF and two Excel files

## ACKNOWLEDGMENTS

We thank Karen Arndt, Stirling Churchman, Alexandra Elchert, and Gyan Prakash for helpful comments on the manuscript. We are grateful to Milan Fábry, Pavel Srb, Veronika Krejčiříková, Magdaléna Hořejší, and Marcela Mádlíkováfor for help with the preparation of NMR samples and data acquisition. We thank Laura McCullough and Tim Formosa for helpful advice and for sharing reagents, and Sanchirmaa Namjilsuren and Karen Arndt for advice on ChIP-seq. We also thank Jon Markert and Lucas Farnung for helpful discussions, and Bobby Hollingsworth for help with protein purification. We also thank the Bauer Core at Harvard University, the Taplin Mass Spectrometry Facility at Harvard Medical School, and the Core for Computational Biomedicine at Harvard Medical School. This work was supported by National Institutes of Health grant R01GM135251 (F.W.) and Czech Science Foundation (GACR) EXPRO grant 25-15442X (V.V.).

## AUTHOR CONTRIBUTIONS

The NMR experiments were conceived of and carried out by V.L. and V.V. The other experiments were conceived of by J.L.W. and F.W., and carried out by J.L.W. The manuscript was written with contributions from all authors.

## DECLARATION OF INTERESTS

The authors declare no competing interests.

## STAR METHODS

### KEY RESOURCES TABLE

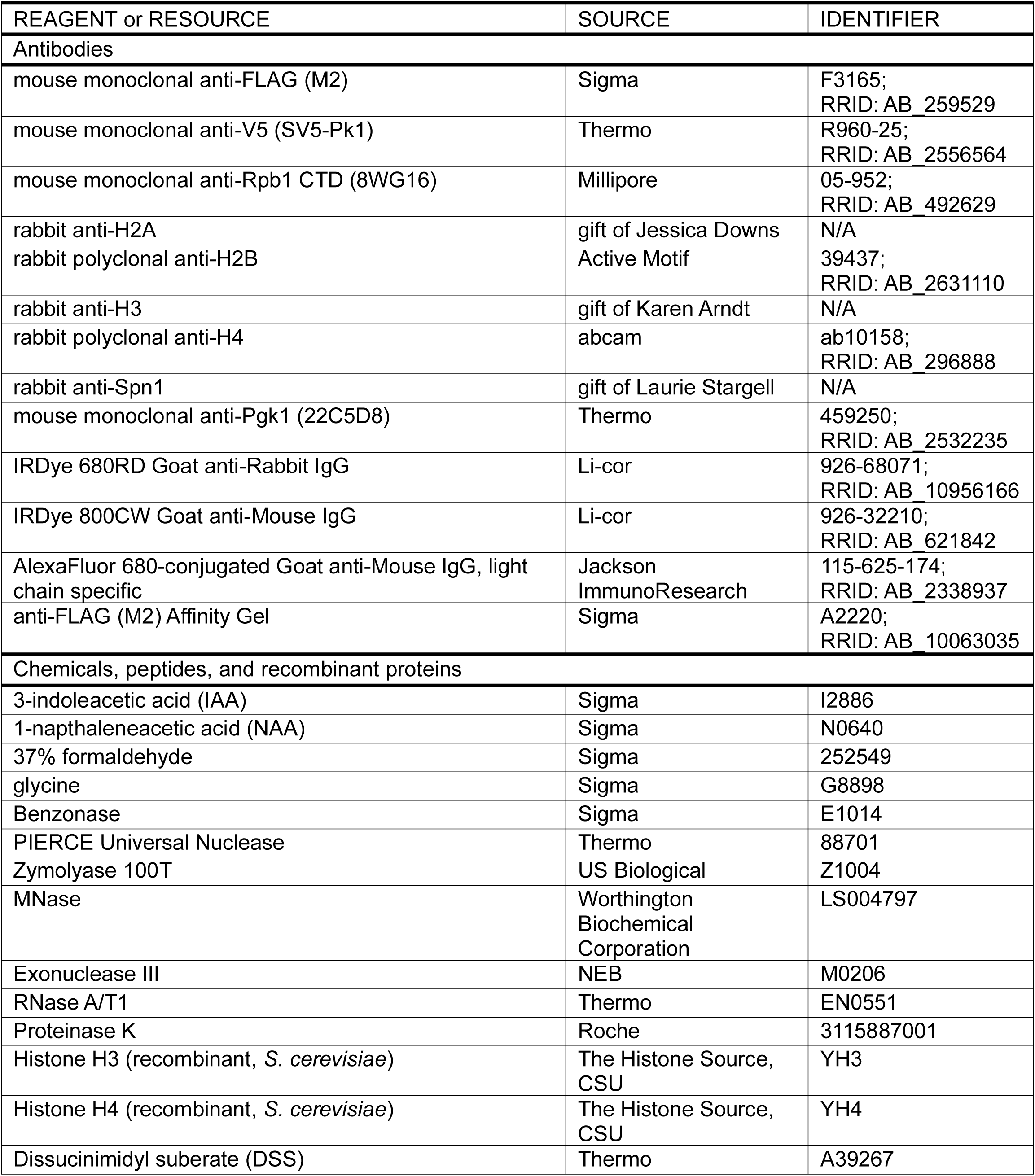
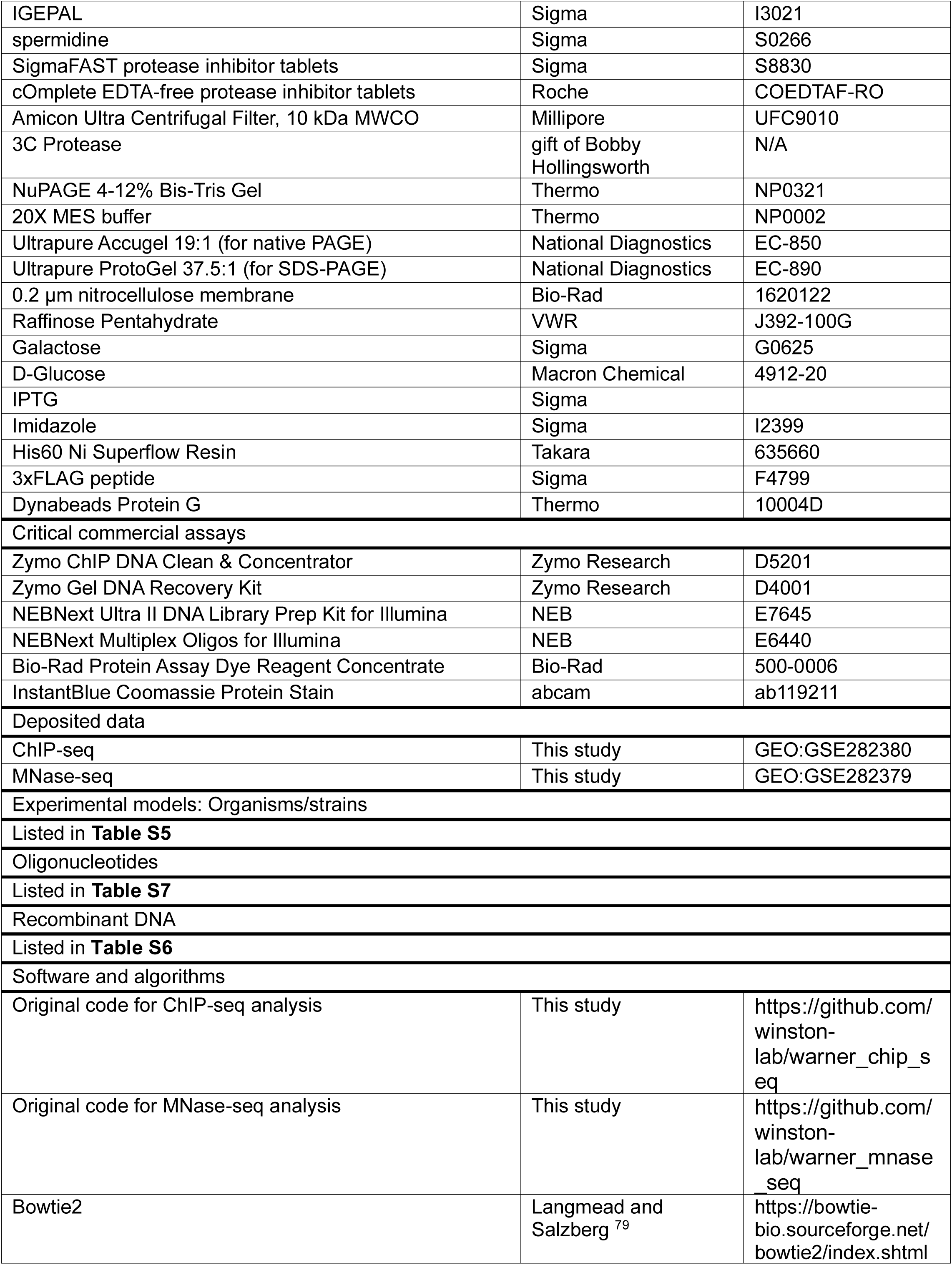
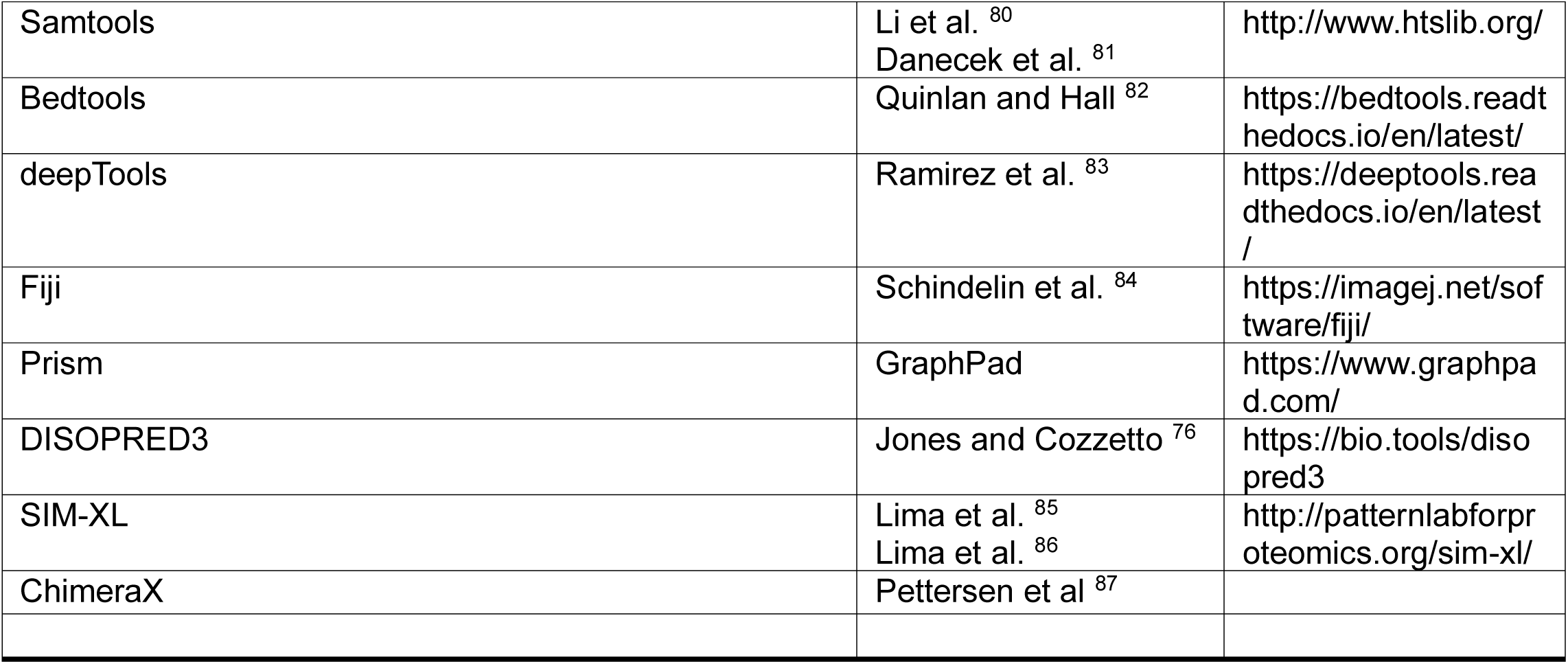

### Yeast strains and growth media

All *S. cerevisiae* strains used in this study (Table S5) are isogenic to S288C^88^. Strains were constructed by standard methods, including transformations, crosses^89^, or CRISPR-Cas9 genome editing^90^. *S. cerevisiae* liquid cultures were grown at 30°C in either YPD (1% yeast extract, 2% peptone, 2% glucose) or synthetic complete media lacking the amino acid of interest (0.2% dropout mix^89^, 0.145% yeast nitrogen base without amino acids and ammonium sulfate, 5% ammonium sulfate, 2% glucose), as indicated. *S. pombe* liquid cultures for spike-in normalization were grown at 32°C in YES (0.5% yeast extract, 3% glucose, 225 mg/l each of adenine, histidine, leucine, uracil, and lysine). Solid media additionally contained 2% agar. Auxin depletion was performed as previously described^23,36^, using 25 μM 3-indoleacetic acid (IAA) for liquid media and 500 μM 1-napthaleneacetic acid (NAA) for solid media.

### Yeast dilution plating (spot tests)

Yeast cultures were grown to saturation overnight from single colonies, normalized by their OD_600_ values, serially diluted 10-fold, and spotted onto the indicated media. Plates were incubated at 30°C unless otherwise noted. All experiments were performed at least in triplicate and a representative example is shown.

### Sequence analysis of Spt6 for charge and disorder

The Spt6 amino acid sequence was analyzed for charge distribution using a custom Python script and plotted using GraphPad Prism. Acidic and basic regions of the Spt6 NTD were identified using the rolling average charge plot followed by manual inspection of the amino acid sequence to define the borders of the regions. Predicted disorder of the Spt6 amino acid sequence was calculated using DISOPRED3^76^ and plotted using GraphPad Prism.

### Construction and analysis of *spt6* mutants

A series of nine mutants was constructed by CRISPR-Cas9 genome editing^90^ in which the charged residues within each of the five acidic or four basic regions of the Spt6 NTD were substituted with alanines. The mutants were tested for growth by spot tests in the following conditions: YPD 30°C (permissive), YPD 37°C (temperature sensitive), and SC-Lys (suppression of *lys2-128δ* (Spt phenotype)^77^). To confirm that the phenotypes were linked to the *spt6* mutation, mutants were crossed with a wild-type *SPT6* strain and tetrads were dissected. All mutants showed single-gene segregation patterns linked to the *spt6* mutations. One allele, *spt6-2001*, displayed variable colony sizes. Sanger sequencing determined that the larger colonies had internal deletions of the acidic-to-alanine mutated region, suggesting that the large number of alanine residues created a neomorphic phenotype. Mutants were subsequently constructed that deleted the acidic regions DE1, DE2, and DE4. These mutants were tested for growth in the same conditions as for the alanine mutants, as well as SC-His (cryptic initiation, *pGAL1-FLO8-HIS3*^41^). Three independent transformants were tested per mutant allele.

### Protein expression and purification

All plasmids used for expression and purification of proteins utilized the phage T7 promoter for a high level of transcription in *E. coli* and are listed in Table S6. Proteins were overexpressed in *Escherichia coli* NEB BL21 (DE3) (C2527H) containing the pRare2 plasmid isolated from Rosetta 2 cells (Novagen 71397-3). For isotopically labelled proteins for NMR studies, bacterial cultures were grown in minimal media containing ^15^N-ammonium sulphate and ^13^C-glucose as the sole nitrogen and carbon sources and 100 μg/ml ampicillin or 50 μg/ml kanamycin, 25 μg/ml chloramphenicol. For other proteins used in NMR experiments (except histones, for histone see below), bacteria were grown in LB medium 100 μg/ml ampicillin or 50 μg/ml kanamycin, 25 μg/ml chloramphenicol, and 0.5% glycerol. For all other experiments, cultures were grown in TY broth supplemented with 50 μg/ml kanamycin and 25 μg/ml chloramphenicol. All cultures were grown at 37°C to OD_600_ ∼ 0.8 and then IPTG (isopropyl-β-D-thiogalactopyranoside) was added to final concentration of 400 μM IPTG. Growth was continued overnight at 18°C. Cells were harvested by centrifugation at 5000g for 20 minutes and then frozen at −20°C before purification. Bacterial pellets were resuspended in lysis buffer (25 mM Tris–HCl pH 7.5, 1 M NaCl, 2 mM β-mercaptoethanol, 10 μM EDTA; 10 ml buffer per 1 gram of pellet) containing protease inhibitors (cOmplete™, EDTA-free, Roche), DNase I (DN25 Sigma-Aldrich, 2 units/mL) and lysed by Emulsiflex C3 (ATA Scientific). In the first purification step, the lysates were cleared by centrifugation (25,000g, 4°C, 30 minutes) and loaded onto HIS-Select^®^ Nickel Affinity Gel resin (Sigma). The bound proteins (except for yeast Spt6 constructs) were eluted by imidazole buffer (25 mM Tris-HCl, 1 M NaCl, 2 mM β-mercaptoethanol, 250 mM imidazole, pH 7.5). The fractions containing purified protein were dialyzed overnight in lysis buffer with TEV protease (for His_6_ tag removal). In the second purification step, the samples were loaded onto HIS-Select^®^ Nickel Affinity Gel resin to remove the TEV protease and proteins nonspecifically interacting with the resin. Yeast Spt6 proteins (1-238 and 1-335) were eluted from HIS-Select^®^ Nickel Affinity Gel resin by the following buffer: 25 mM Tris-HCl, 250 mM NaCl, 2 mM β-mercaptoethanol, 250 mM imidazole, pH 7.5. For proteins for NMR studies, fractions were then dialysed against 25 mM Tris-HCl, 150 mM NaCl, 2 mM β-mercaptoethanol, pH 7.5 in the presence of GST-tagged preScission protease. The following day, the dialysed samples were passed through a GST column, to remove the preScission protease and through HIS-Select^®^ Nickel Affinity Gel resin to remove the His-MBP-tag. The purified proteins present in the flow-through fraction of HIS-Select^®^ Nickel Affinity gel resin were concentrated using a Vivaspin centrifugal filtration unit with 10 kDa cutoff. For all other experiments, imidazole eluted fractions were cleaved with 3C protease (kind gift of Bobby Hollingsworth) overnight at 4°C. The cleaved samples were bound to anti-Flag M2 resin (Sigma) to remove the His-MBP-tag and the protease and eluted with the following buffer: 50 mM HEPES pH 7.5, 300 mM NaCl, 100 μg/ml 3xFlag peptide. The purified proteins were concentrated using an Amicon Ultra centrifugal filtration unit with a 10 kDa cutoff. In all cases, the final purification step was size exclusion chromatography using Superdex 75 or 200 10/300 GL in a buffer that contained 25 mM Tris-HCl, 150 mM NaCl, 1 mM TCEP, and 10 μM EDTA pH 7.5. The purest fractions were concentrated as before. The final concentration of the purified protein was determined on a NanoDrop spectrophotometer and its purity was evaluated by SDS-PAGE followed by Coomassie Brilliant Blue. The purified proteins were frozen in liquid nitrogen and stored at −80°C.

*Xenopus laevis* histone proteins (H2A, H2B, H3, H4) were expressed in *E. coli* BL21 CodonPlus (DE3) RIL cells (Agilent technologies 230245) using a YT medium 50 μg/ml ampicillin and 25 μg/ml chloramphenicol and 1% glucose as an insoluble fraction, as described earlier^1,91^. To prepare isotopically labeled histones, they were expressed in minimal medium containing ^15^N-ammonium sulphate and ^13^C-glucose as the sole nitrogen and carbon sources and 50 μg/ml ampicillin and 25 μg/ml chloramphenicol. All cultures were grown at 37°C until induction of protein expression at OD_600_ ∼ 0.5 with 400 μM IPTG (isopropyl-β-D-thiogalactopyranoside). Protein expression into inclusion bodies continued 6h at 37°C. All histone variants were prepared the same as previously described^92^. Briefly, after extraction from the inclusion bodies, the histones were purified on an anion exchange column (HiPrep^®^ Q FF 16/10, GE Healthcare) and a cation exchange column (HiPrep^®^ SP FF 16/10, GE Healthcare) using anion (7 M deionized urea, 20 mM Tris–HCl, 0.2-1 M NaCl, 1 mM EDTA, 5 mM β-mercaptoethanol, pH 7.5) and cation (7 M deionized urea, 20 mM Na-acetate, 0.1-1 M NaCl, 1 mM EDTA, 5 mM β-mercaptoethanol, pH 5.2) exchange buffers. Pure fractions were pooled and dialyzed against 2 mM β-mercaptoethanol in MilliQ water, flash-frozen, lyophilized and stored at −80 °C.

*X. laevis* histone H2A/H2B dimers or H3/H4 dimers/tetramers were refolded as previously described for octamers^1,91^. Briefly, equimolar amounts of individual histones were resuspended in unfolding buffer (6 M GuHCl, 20 mM Tris–HCl, 5 mM DTT, pH 7.5), mixed in a final concentration of 1 mg/ml, and refolded during dialysis into 10 mM Tris–HCl, 2 M NaCl, 1 mM EDTA, 5 mM β-mercaptoethanol, pH 7.5 (repeated 3X). The samples were concentrated and applied to a Superdex^®^ 200 Increase 10/300 GL column (GE Healthcare) that was equilibrated in 25 mM Tris-HCl, 200 mM NaCl, 10 µM EDTA and 2 mM TCEP buffer pH 7.5. Histone dimer/tetramer containing fractions were pooled, concentrated, flash frozen and stored at −80°C. The samples used for NMR spectra of H2A/H2B dimers were prepared using either ^13^C/^15^N labeled H2A with natural abundance H2B or ^13^C/^15^N labeled H2B with natural abundance H2A, respectively.

*S. cerevisiae* histones H2A and H2B were co-expressed in *E. coli* and purified as a complex as described previously^93^. Briefly, cell lysates were prepared as described above for Spt6 NTD constructs, except that the buffer contained 2M NaCl. Proteins were purified using His60 Ni Superflow Resin (Takara), concentrated, and applied to a Superdex^®^ 200 Increase 10/300 GL column (GE Healthcare). The fractions containing H2A/H2B dimers were collected, pooled, concentrated, flash frozen, and stored at −70°C. *S. cerevisiae* histones H3 and H4 were purchased from The Histone Source (Colorado State University) and refolded as described above.

### Electrophoretic mobility shift assays

Electrophoretic mobility shift assays (EMSAs) were performed as previously described^50,94^, with minor modifications. Individual reactions were 20 μl in volume. Purified Spt6 fragments and yeast histones were added to the indicated final concentrations in a buffer that contained 20 mM HEPES pH 7.5, 300 mM NaCl, 1 mM EDTA, 1 mM β-mercaptoethanol, 2% glycerol (v/v), and 12% sucrose (w/v). Reactions were incubated at 30°C for 10 minutes. Half of the reaction (10 μl) was separated by native polyacrylamide gel electrophoresis (PAGE) for one hour at 80 volts on a 0.4X TBE acrylamide (5%, 19:1, National Diagnostics) gel. The other half was mixed with an equal volume of 2X Laemmli buffer^95^ and separated under denaturing conditions on a Tris-Glycine acrylamide (15%, 37.5:1, National Diagnostics) gel^95^. To visualize proteins, gels were stained with InstantBlue Coomassie Protein Stain (Abcam) according to the manufacturer’s specifications. The gels were imaged on a Licor Aerius, using the 700 nm channel, and the images were processed using ImageJ.

### In vitro crosslinking and analysis

Purified Spt6(1-238) and yeast histones (either H2A/H2B or H3/H4) were mixed to a final concentration of 5 μM each in a buffer that contained 20 mM HEPES pH 7.5, 300 mM NaCl, 1 mM β-mercaptoethanol, and 2% glycerol (v/v). Reactions were incubated at 25°C for 15 minutes. Discuccinimidyl suberate (DSS, Thermo) in DMSO was then added to the reactions at a final concentration of 2.5 mM and crosslinking was allowed to proceed for 30 minutes at 25°C. Tris-HCl pH 7.5 was added to a final concentration of 50 mM and the reactions were incubated at 25°C for 15 minutes to quench any unreacted crosslinker. Samples analyzed by SDS-PAGE were mixed with 2X Laemmli buffer, separated on 15% acrylamide Tris-Glycine gels, and visualized by staining with InstantBlue Coomassie Protein Stain (Abcam). For samples analyzed by mass spectrometry, a fraction of the reaction was analyzed by SDS-PAGE to confirm crosslinking and the remainder was submitted to the Taplin Mass Spectometry Facility at Harvard Medical School for LC/MS/MS analysis. LC/MS/MS results were analyzed and visualized using Spectrum Identification Machine for Cross-Linked Peptides (SIM-XL,^85,86^).

### Peptide synthesis

Spt6 derived peptides were synthesized by solid phase synthesis in the Laboratory of Medicinal chemistry, IOCB, ASCR v. v. i., Prague, Czech Republic.

### NMR spectroscopy

NMR spectra were acquired at 25°C on an 850 MHz Bruker Avance spectrometer using a triple-resonance (^15^N/^13^C/^1^H) cryoprobe. The samples for assignments of the Spt6, Spn1 and Elf1 constructs were prepared at 0.3-0.5 mM in a volume of 0.35 mL, in buffer (25 mM Tris-HCl pH 7.5, 200 mM NaCl, 1 mM TCEP, 10 µM EDTA,), 5 % D_2_O/95% H_2_O. Samples that were titrated by H3/H4 histone dimers/tetramers were measured in buffer with 300 mM NaCl. A series of double- and triple-resonance spectra^96,97^ were recorded to obtain sequence-specific backbone resonance assignments for the Spt6, Spn1 and Elf1 constructs. The H2A/H2B samples, where one of the histones was isotopically labelled, were measured in 25 mM MES pH 6.0, 400 mM NaCl, consistent with conditions used for a published backbone resonance assignment^98^.

To monitor changes upon protein binding, we calculated either the free-to-bound signal ratio of individual peaks in the 3D HNCO spectra or the chemical shift perturbations (CSPs) in the 2D ^15^N/^1^H HMQC spectra. The NMR spectra were analysed using NMRFAM-SPARKY^99^, and the signal ratio or CSP graphs were plotted using GraphPad Prism. Signal ratios were used for determination of all binding profiles for Spt6, Spn1 and Elf1 constructs, while CSPs were employed to characterize the binding of Spt6-derived peptides to H2A/H2B. The CSP of each assigned resonance in the 2D ^15^N/^1^H HMQC spectra of the H2A or H2B was calculated as the geometrical distance (in ppm) between the peak in the 2D ^15^N/^1^H HMQC spectra acquired before and after peptide addition using the formula: 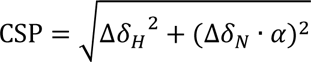, where *α* is a weighing factor of 0.2 used to account for differences in the proton and nitrogen spectral widths^100^. In binding experiments the 50 μM ^13^C/^15^N-labeled proteins were measured in the presence of equimolar unlabelled proteins or peptides and compared with DMSO (in the cases of peptides) or a buffer control. For each interaction 1D ^1^H spectra, the ^15^N/^1^H HMQC spectra in the SOFAST fashion^101^ and 3D HNCO spectra were recorded.

### Co-immunoprecipitation

*S. cerevisiae* cells with plasmids expressing full-length or Spt6Δ2-238 constructs were grown in SC-Trp to OD_600_ ≈ 0.6 at 30°C. The cells were pelleted, washed once in PBS, and frozen at −70°C. *S. cerevisiae* cells with plasmids expressing Spt6 NTD constructs were grown in SC-Ura with 2% raffinose to OD_600_ ≈ 0.6 at 30°C. Transcription of the *SPT6* sequence was induced by the addition of galactose to a final concentration of 2% for one hour. The cells were pelleted, washed once in PBS, and frozen at −70°C. Frozen pellets were resuspended in 500 μl of IP Buffer (100 mM HEPES-KOH pH 7.9, 100 mM KOAc, 10 mM Mg(OAc)2, 2 mM EDTA, 0.4% NP-40/IGEPAL, 1mM DTT, 1X SigmaFast EDTA-free Protease Inhibitor, modified from^58^). Approximately 300 μl of acid-washed glass beads were added and the cells were lysed by bead beating at 4°C for 7 minutes with a 4-minute incubation on ice after every minute of bead beating. The resulting crude extract was split into two 250 μl aliquots and frozen at −70°C. Aliquots were thawed on ice and 1 μl of Benzonase nuclease (Sigma) or Universal nuclease (Thermo) was added prior to incubation for 30 minutes at 4°C on a rotisserie. The tubes were then centrifuged at 12,500 rpm for 10 minutes and the supernatant was removed to a new tube. Protein concentrations were determined by Bradford assay (Bio-rad) and normalized across IPs with IP buffer to a final volume of 230 μl. 12 μl of each reaction was taken as input and mixed with 2X Laemmli buffer, followed by the addition of 10 μl of anti-FLAG beads (20 μl slurry, Sigma) equilibrated in IP buffer, and incubation for two hours at 4°C on a rotisserie. The beads were washed twice with IP buffer and the bound proteins were eluted by boiling for 10 minutes in 45 μl 1X Laemmli buffer. The samples were separated by SDS-PAGE on 4-12% NuPAGE Bis-Tris gels (Invitrogen), transferred to 0.2 μm nitrocellulose membranes (Bio-rad), and analyzed by Western blotting. The following antibodies were used for Western blots: 1:10000 anti-FLAG (M2, Sigma), 1:6000 anti-Spn1 (generously provided by Laurie Stargell), 1:4000 anti-H2A (generously provided by Jessica Downs), 1:4000 anti-H2B (ActiveMotif), 1:15000 anti-H3 (generously provided by Karen Arndt), 1:1000 anti-H4 (abcam). Western blots were visualized on a Li-cor Aerius Imager using IRDye 680RD Goat anti-Rabbit IgG (Li-cor), IRDye 800CW Goat anti-Mouse IgG (Li-cor), or AlexaFluor 680-conjugated Goat anti-Mouse IgG, light chain specific (Jackson ImmunoResearch). Blot images were quantified using ImageJ and the results were plotted using GraphPad Prism.

### Chromatin immunoprecipitation and sequencing (ChIP-seq)

Strain FY3544 with plasmids pJLW93 or pJLW95 was inoculated into SC-Trp from a single colony and grown at 30°C to OD_600_ ≈ 0.6. The cultures were split and diluted in SC-Trp to OD_600_ = 0.3 and then treated with either DMSO (vehicle) or 3-indoleacetic acid (depletion). After 90 minutes of growth at 30°C, the cultures were brought to 25°C by mixing with 4°C media and crosslinked with formaldehyde (1% final concentration) for 30 minutes with shaking at room temperature. Glycine was added to quench the crosslinking reactions at a final concentration of 125 mM and shaking was continued for 10 minutes. The cells were then pelleted, washed once in cold TBS, and frozen at −70°C. Expression of plasmid-encoded Spt6 and depletion of genome-encoded Spt6 was confirmed by western blot (Figure S5F).

Cell pellets were resuspended in LB140 (50 mM HEPES-KOH pH7.4, 140 mM NaCl, 1 mM EDTA, 1 % TritonX-100, 0.1% sodium deoxycholate, 0.2% SDS, 1X SigmaFAST protease inhibitor cocktail (Sigma), 1 mM PMSF, 0.4 mM DTT) and lysed by bead beating at 4°C for 8 minutes, with 4 minutes on ice in between every minute of bead beating. Chromatin was isolated by centrifugation at 12,500 rpm at 4°C for 30 minutes in an Eppendorf centrifuge, washed once with LB140, and repelleted. Chromatin was resuspended in 1 ml of LB140 and sonicated on a Covaris sonicator (450 seconds per sample, 140 peak power, 5.0 duty factor, 200 cycles per burst) to fragments smaller than 1 kb with most fragments smaller than 500 bp as determined by agarose gel electrophoresis. Protein concentration was measured by Bradford assay (Biorad). 400 μg of chromatin was used per IP. Samples were brought to the same concentration with LB140, *S. pombe* chromatin was added at 10% (w/w) as a spike-in, and IP reactions were diluted 1:2 in WB140 (50 mM HEPES-KOH pH 7.4, 140 mM NaCl, 1 mM EDTA, 1% TritonX-100, 0.1% sodium deoxycholate) to increase IP efficiency. A 5% equivalent of one IP reaction was saved as input. Antibodies were added to each reaction as follows: 5 μl ɑ-FLAG (M2, Sigma), 5 μl ɑ-V5 (Invitrogen), 10 μl ɑ-Rpb1 (8WG16, EMD Millipore).

Immunoprecipitations were incubated overnight (14-16 hours) at 4°C with end-over-end rotation. 50 μl of Protein G Dynabeads (Thermo) equilibrated in WB140 were added to the IPs and samples were incubated at 4°C for 5 hours with end-over-end rotation. Beads were washed twice each with WB140, WB500 (50 mM HEPES-KOH pH 7.4, 500 mM NaCl, 1 mM EDTA, 1% TritonX-100, 0.1% sodium deoxycholate), and WBLiCl (50 mM Tris-HCl pH 7.5, 250 mM LiCl, 1 mM EDTA, 0.5% IGEPAL, 0.5% sodium deoxycholate) and once with TE (10 mM Tris-HCl ph 7.5, 1 mM EDTA). The immunoprecipitated material was eluted by addition of 100 μl TES (50 mM Tris-HCl pH 7.5, 10 mM EDTA, 1% SDS) and incubation at 65°C for 15 minutes. Elutions were performed twice and pooled. Eluates were incubated at 65°C overnight to reverse crosslinking. 200 μl of TE was added to eluates. 8 μg of RNase A/T1 (Thermo) was added and the reactions were incubated at 37°C for 2 hours. 160 μg of Proteinase K (Roche) was added anf the reactions were incubated at 42°C for 2 hours. DNA was purified using a Zymo ChIP DNA Clean & Concentrate kit following manufacturer specifications. The purified DNA was used to build sequencing libraries using the NEBNext Ultra II DNA library Prep Kit for Illumina (NEB) following manufacturer specifications. Libraries were indexed using NEBNext Multiplex Oligos for Illumina (NEB). Libraries were sequenced at the Bauer Core Facility at Harvard University on a NovaSeq 6000.

### ChIP-seq analysis

Sequencing results were aligned to the *S.cerevisiae* and *S. pombe* (spike-in) genomes using Bowtie2^79^ and sorted using samtools^80,81^. Spike-in normalization factors were calculated using a custom python script (see Data and code availability). Spike-in normalized coverage tracks were generated using deeptools^83^. Pearson correlation coefficients between all libraries were also calculated using deeptools. ChIP experiments were performed in biological triplicate. Overall, replicates correlated well with one another (Figure S5A). Correlation plots, heatmaps, and metagene plots were generated using a custom python script.

### Isolation and genetic analysis of suppressor mutations

Yeast strain FY3552 was used to isolate spontaneous suppressors of the *spt6ΔDE2DE4* temperature-sensitive phenotype. Cultures were inoculated from five single colonies and grown overnight in liquid YPD. Approximately 1 x 10^7^ cells from each culture were spread on YPD plates and the plates were incubated at 37°C. Four plates yielded a small number of colonies (four to eight) and one plate produced only a single colony. The single colony plus two colonies from each of the other four plates were purified, yielding nine suppressor candidates. The purified candidates were retested for growth at 37°C. To test for dominance, the suppressor strains were crossed with a *spt6ΔDE2DE4* strain (FY3555) and the resulting diploids were tested for growth at 37°C. The diploids were sporulated and the resulting tetrads were dissected to analyze segregation of the suppression phenotype. To test for complementation, the suppressors were crossed to each other and the resulting diploids were tested for growth at 37°C. These genetic analyses resulted in the identification of six independent suppressor mutants in two complementation groups.

### Identification of suppressor mutations

Genomic DNA was extracted from one suppressor mutant from each complementation group (FY3553 and FY3554) and the parent (FY3552) using standard protocols^102^. Library preparation and sequencing was performed by Seqcenter (Pittsburgh, PA). To identify candidate suppressor mutations, each suppressor genome was compared to the parent using MutantHuntWGS^103^. Variants that mapped within open reading frames in each suppressor genome were manually inspected and candidate mutations in *SPT16* and *POB3* were identified. Similar mutations were identified by Sanger sequencing the same genes in the other four suppressor mutants, with one complementation group representing mutations in *SPT16* and the other representing mutations in *POB3*. To confirm that suppression was caused by mutations in *SPT16* and *POB3*, the suppressor mutations were transformed by plasmids that contain the wild-type *SPT16* and *POB3* genes and shown to be complemented by the expected plasmid based on the mutant sequence.

### Micrococcal nuclease digestion and sequencing (MNase-seq)

Strains FY3544 and FY3546 with plasmid pRS414, pJLW93, or pJLW95 were inoculated into SC-Trp from a single colony and grown at 30°C to OD_600_ ≈ 0.6. The cultures were split and diluted in SC-Trp to OD_600_ = 0.3 and then treated with either DMSO (vehicle) or 3-indoleacetic acid (depletion). After 90 minutes of growth at 30°C, the cells were pelleted, washed once in cold TBS, and frozen at −70°C. Cell pellets were processed for MNase digestion as described previously^104^. Briefly, cell pellets were resuspended in water and crosslinked with 1% formaldehyde for 15 minutes at room temperature. Crosslinking was quenched by the addition of glycine (125 mM final) and cells were pelleted by centrifugation. Cells were resuspended in spheroplasting buffer (125 μg/ml Zymolyase 100T, 1 M sorbitol, 50 mM Tris-HCl pH 7.5, 5 mM β-mercaptoethanol) and incubated at room temperature with end-over-end rotation until >90% of the cells were spheroplasts as assessed by microscopy. The spheroplasts were collected and resuspended in MNase digestion buffer (1 M sorbitol, 50 mM NaCl, 10 mM Tris-HCl pH 7.5, 5 mM MgCl_2_, 1 mM CaCl_2_, 0.074% NP-40, 500 μM spermidine, 1 mM β-mercaptoethanol). An amount of MNase (Worthington) optimized to produce mononucleosomes (between 100 and 160 U) and 30 U of Exonuclease III (NEB) was added to each sample and the reactions were incubated at 37°C for exactly 15 minutes. Digestion was halted by the addition of STOP buffer (50 mM EDTA, 50 mM EGTA) and purified *S. pombe* mononucleosomal DNA was spiked in at a ratio of 1 ng *S. pombe* DNA / 1.5 x 10^7^ *S. cerevisiae* cells. Samples were digested with RNase A/T1 (Thermo) and Proteinase K (Roche) and crosslinks were reversed by addition of SDS (1% final) and incubation at 65°C for 45 minutes. DNA was purified using ChIP DNA Clean & Concentrator columns (Zymo). DNA was separated by agarose gel electrophoresis and the band representing the mononucleosomal fragment was excised and DNA was isolated using a Zymoclean Gel DNA Recovery Kit (Zymo). 50 ng of DNA was used to build sequencing libraries using the NEBNext Ultra II DNA library Prep Kit for Illumina (NEB) following manufacturer specifications. Libraries were indexed using NEBNext Multiplex Oligos for Illumina (NEB). Libraries were sequenced at the Bauer Core Facility at Harvard University on a NovaSeq 6000.

### MNase-seq analysis

Sequencing results were aligned to the *S.cerevisiae* and *S. pombe* (spike-in) genomes using Bowtie2^79^ and sorted using samtools^80,81^. Spike-in normalization factors were calculated using a custom python script (see Data and code availability). Spike-in normalized coverage tracks of nucleosome dyad positions were generated using deeptools^83^. Pearson correlation coefficients between all libraries were also calculated using deeptools. MNase experiments were performed in biological triplicate. Overall, replicates correlated well with one another (Figure S6A). Correlation plots, heatmaps, and metagene plots were generated using a custom python script.

### Data and code availability

All next-generation sequencing data (unprocessed read files and processed coverage files) can be accessed at GEO (https://www.ncbi.nlm.nih.gov/geo/) with accession numbers GSE282379 (MNase-seq) and GSE282380 (ChIP-seq). All scripts used to analyze next-generation sequencing data and an explanation of how spike-in normalization factors were calculated can be accessed at https://github.com/winston-lab.

## Notes

### Competing Interest Statement

The authors have declared no competing interest.

