## supplemental figures and tables for "The histone chaperone Spt6 controls chromatin structure through its conserved N-terminal domain"

This supplemental information contains:

**Figure S1** - related to Figure 1. The acidic NTD is a conserved feature of Spt6

**Figure S2** - related to Figure 2. In vitro analysis of Spt6 NTD-histone binding.

**Figure S3** - related to Figure 3. NMR analysis of histones H2A and H2B interactions with Spt6 NTD peptides.

**Figure S4** - related to Figure 3. NMR analysis of the Spt6 NTD, Spn1, and Elf1 uncovers complex regulation of histone binding activities.

**Figure S5** - related to Figure 5. Biological replicates of Spt6 and Rpb1 ChIP-seq experiments.

**Figure S6** - related to Figure 6. The Spt6 NTD and FACT cooperate to control chromatin structure.

**Figure S7** - related to Figure 7. Structural models supporting the cooperation between Spt6 and FACT in co-transcriptional histone transfer.

**Table S1** – alanine scan phenotypes

**Table S2** – mass spec results; not in this pdf, included as an Excel file

**Table S3** – mutant phenotypes

**Table S4** – *spt16* and *pob3* suppressor mutations

**Table S5** – yeast strains

**Table S6** – plasmids

**Table S7** – oligos; not in this pdf, included as an Excel file

Figure S1, related to Figure 1

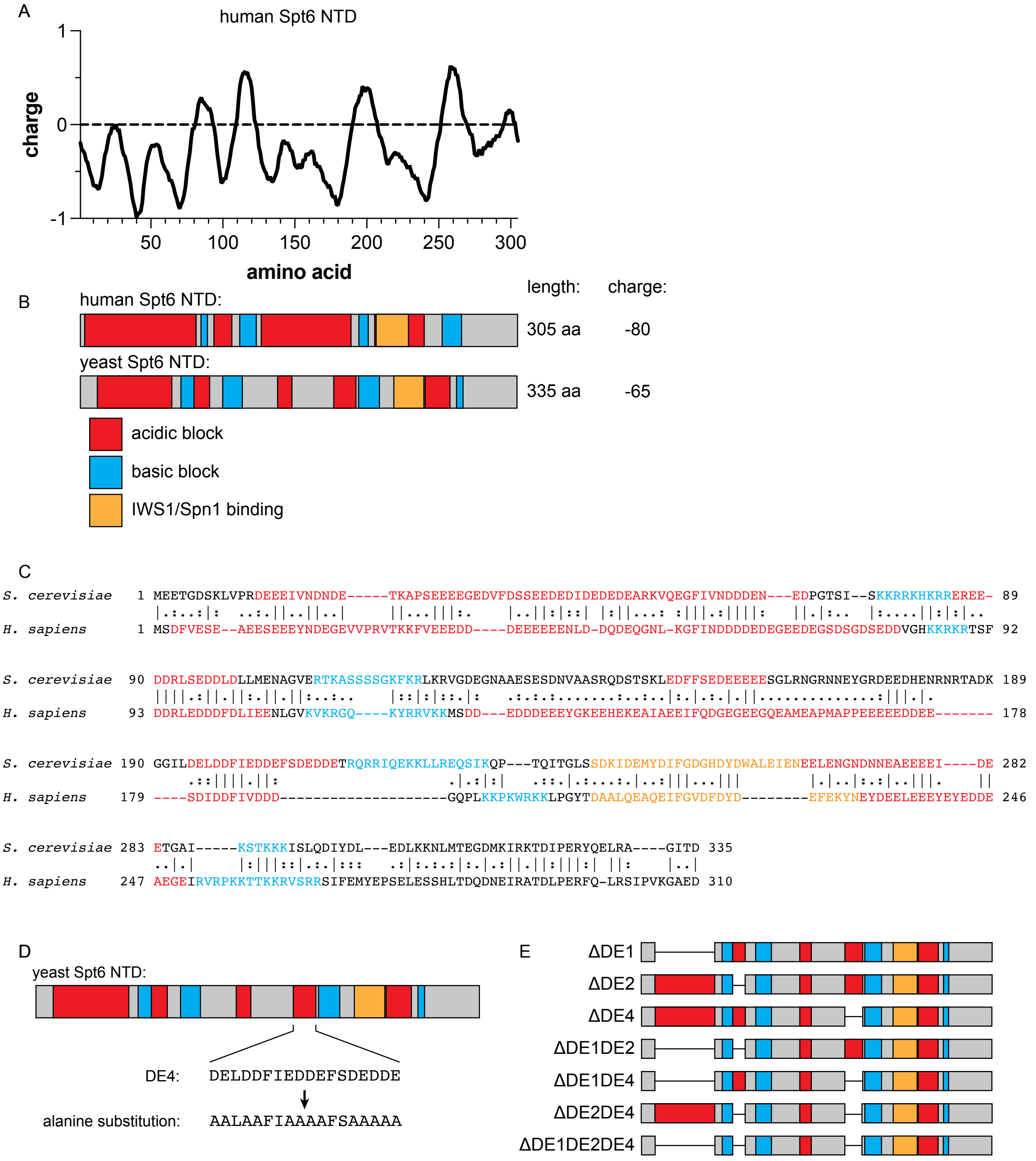


**Figure S1, related to Figure 1. The acidic NTD is a conserved feature of Spt6.**
(A) The human Spt6 NTD is acidic. A line plot showing the rolling average of amino acid side-chain charge of human Spt6 in a fifteen-residue window.
(B) Diagrams of homologous NTDs from human and *S. cerevisiae* Spt6. Acidic (red), basic (blue), and IWS1/Spn1 binding (orange) regions identified by sequence analysis are indicated. The position of the human diagram is aligned with the x-axis of the plot in (A). Numbers to the right indicate the C-terminal end of each domain.

(C) Pairwise amino acid sequence alignments of human and yeast Spt6 NTDs. Residues are colored as in (B) to indicate if they are in acidic, basic, or IWS1/Spn1 binding regions. Alignments were generated using EMBOSS Needle[1]. In the shown alignment, 109/425 (25.6%) of residues are identical and 172/425 (40.5%) of residues are similar.
(D) Schematic illustrating an example of an alanine substitution mutant, in this case a substitution of DE4. All acidic residues within DE4 were substituted with alanine.
(E) Acidic block deletion alleles.

Figure S2, related to Figure 2

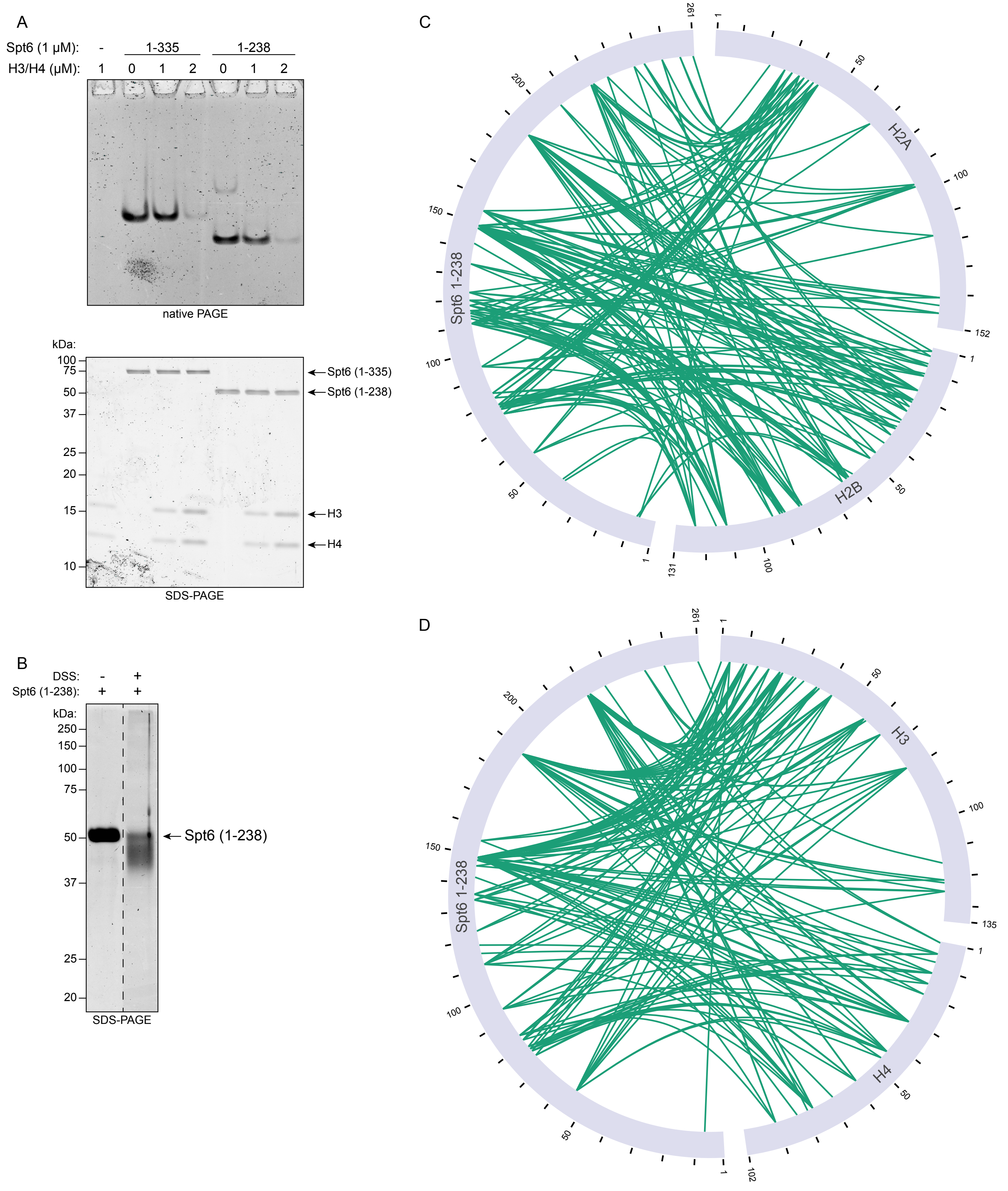

**Figure S2, related to Figure 2. In vitro analysis of Spt6 NTD-histone binding.**
(A) Electrophoretic mobility shift assay (EMSA) analysis of Spt6 NTD-H3/H4 reactions. Spt6 constructs and histones H3/H4 were mixed in the amounts indicated and separated by native of denaturing PAGE. Gels were stained with Coomassie to visualize proteins. Spt6 NTD-H3/H4 complexes are not predicted to enter the gel under native electrophoresis conditions due to their positive charge.
(B) Spt6 (1-238) does not form higher molecular weight crosslinked complexes alone. Purified components were mixed as indicated prior to separation by denaturing PAGE. Gels were stained with Coomassie to visualize proteins. Intervening lanes have been removed for visualization purposes, as marked by the dashed line.
(C and D) Visualization of crosslinks detected by mass spectrometry between Spt6 (1-238) and H2A/H2B (C) and H3/H4 (D). Residue numbering is marked on the outer ring of the arcs depicting the proteins.

Figure S3, related to Figure 3

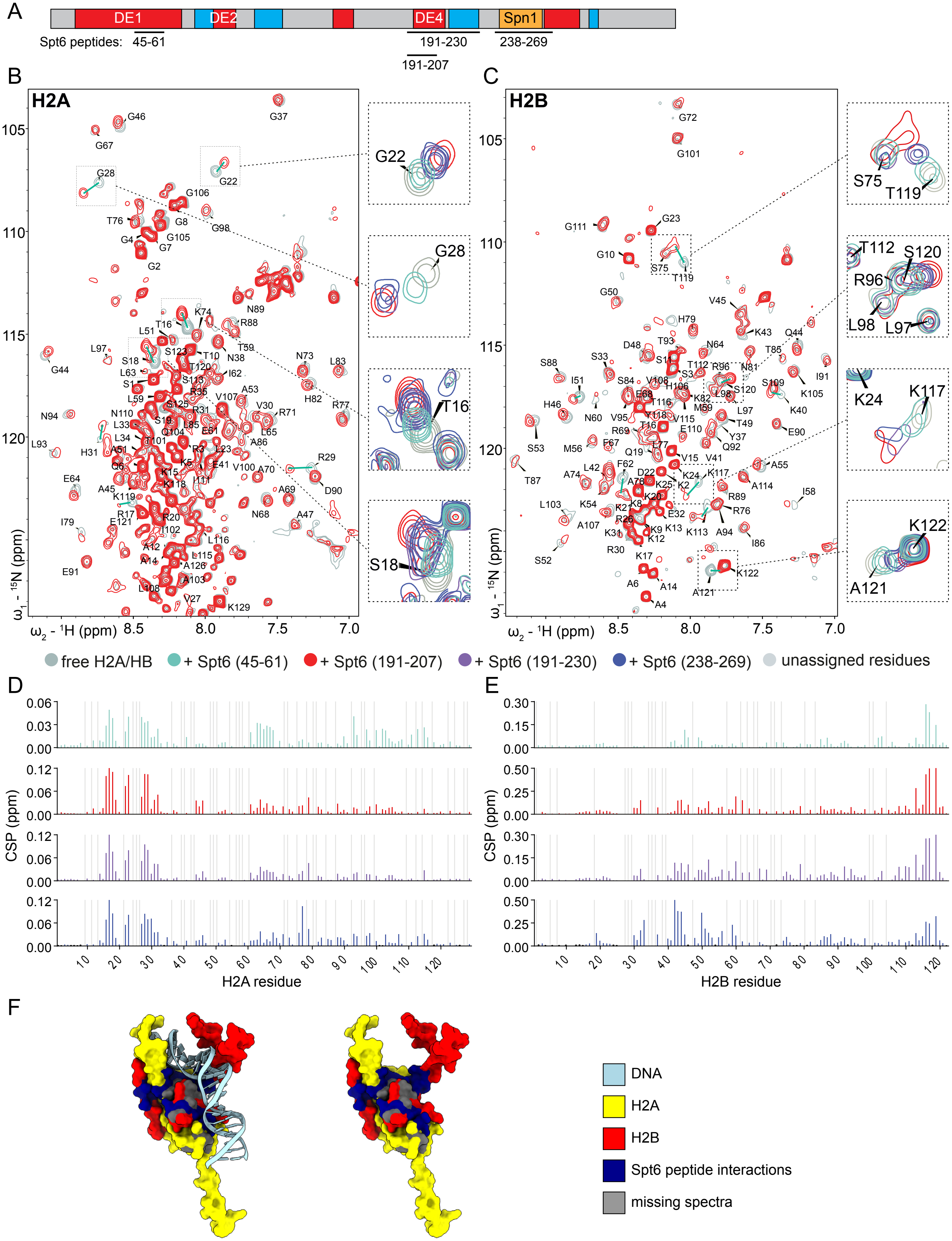

**Figure S3, related to Figure 3. NMR analysis of histones H2A and H2B interactions with Spt6 NTD peptides.**

(A) Schematic diagram of the Spt6 NTD showing acidic blocks (red), basic blocks (blue), and the Spn1-binding region (orange). The Spt6 peptides are indicated below as black lines along with the amino acids contained in each peptide.

(B and C) NMR analysis of the interaction between the Spt6 NTD-derived peptides and an H2A/H2B dimer assembled using either $^{15}N$ labeled H2A (B) or $^{15}N$ labeled H2B (C). Above is the full 2D $^{15}N/^{1}H$ HSQC spectrum obtained using specifically labeled H2A or H2B within the dimer, shown in the absence (gray) or presence of Spt6 peptides (colors). Significant shifts are highlighted by green arrows. Zoomed-in regions show overlays of the spectra obtained after addition of the four different Spt6 peptides.

(D and E) Plots of chemical shift perturbation, summarizing the changes in histone signal positions upon peptide binding. Residues that could not be assigned are shown in gray. NMR analysis reveals that all tested Spt6 NTD-derived peptides bind to a DNA-binding surface of the H2A/H2B dimer.

(F) Spt6 peptides bind to the DNA binding surface of H2A/H2B. Histones H2A/H2B bound to DNA as part of a nucleosome[2] (PDB: 1KX5). Histone H2A is colored yellow, histone H2B is colored red, and DNA is colored light blue. Histone H2A and H2B residues with strong chemical shifts are colored dark blue. Unassigned residues are colored gray.

Figure S4, related to Figure 3

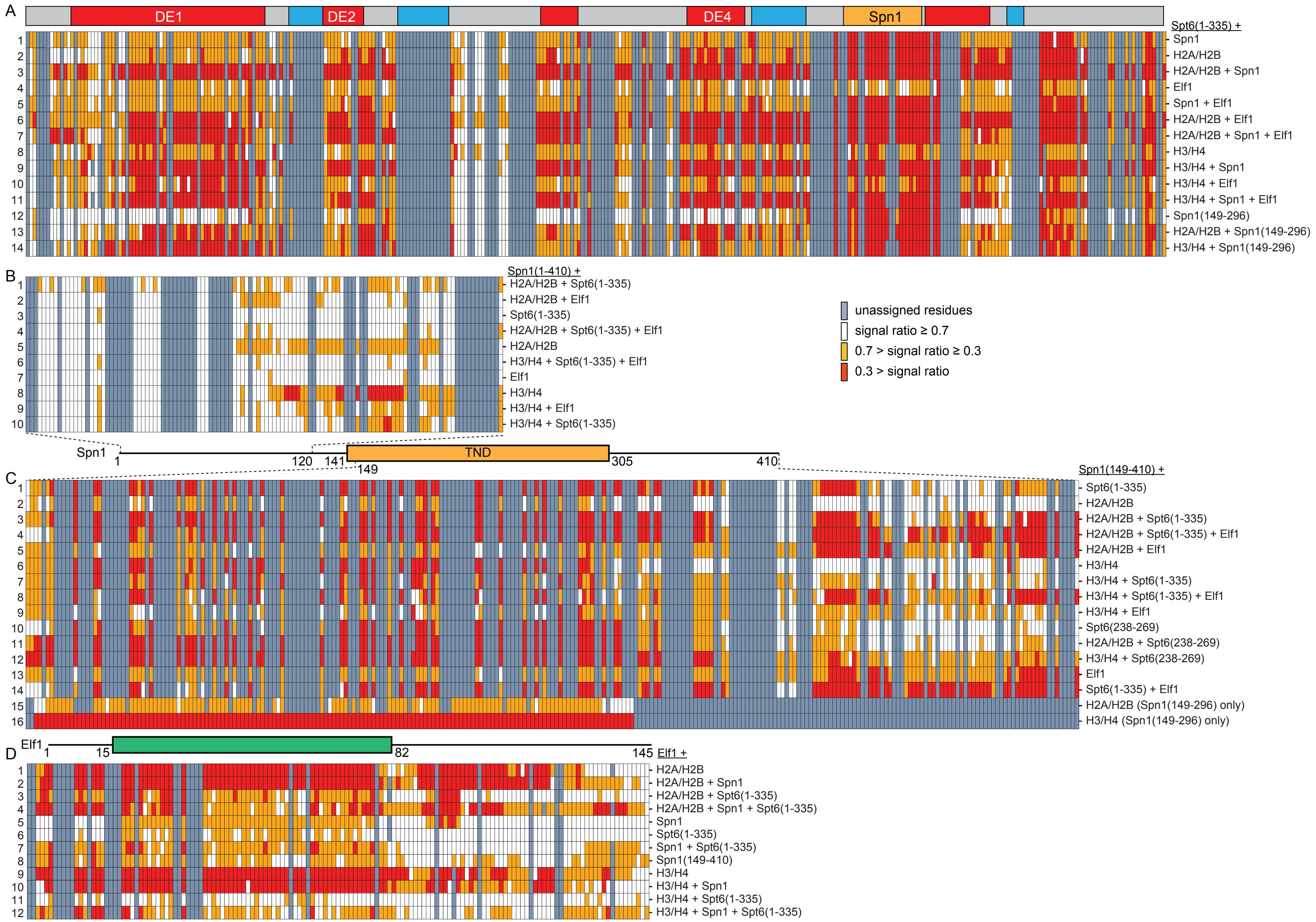

**Figure S4, related to Figure 3. NMR analysis of the Spt6 NTD, Spn1, and Elf1 uncovers complex regulation of histone binding activities.**
(A) Heatmap representations of NMR spectra signal ratios for labeled Spt6 (1-335). Each row of the heatmap represents the results of the Spt6 construct incubated with the indicated binding partners. Above is a schematic illustrating the acidic (red) and basic (blue) regions of the Spt6 NTD, as well as the Spn1 binding region (orange). Below is a key for the heatmap values.
(B) Heatmap representations of NMR spectra signal ratios of the labeled full-length Spn1 construct incubated with the indicated binding partners. Below the heatmap is a schematic of the Spn1 protein with the central core (TND, TFIIS N-terminal Domain-like) labeled. Dotted lines indicate the boundaries of the residues resolved in the NMR experiments.
(C) As in (B), but for Spn1 (149-410).
(D) Above is a schematic illustrating the Elf1 protein with the conserved transcription elongation factor 1 superfamily domain shown as a box. Below are heatmap representations of NMR spectra signal ratios of labeled full-length Elf1 incubated with the indicated binding partners.

Figure S5, related to Figure 5

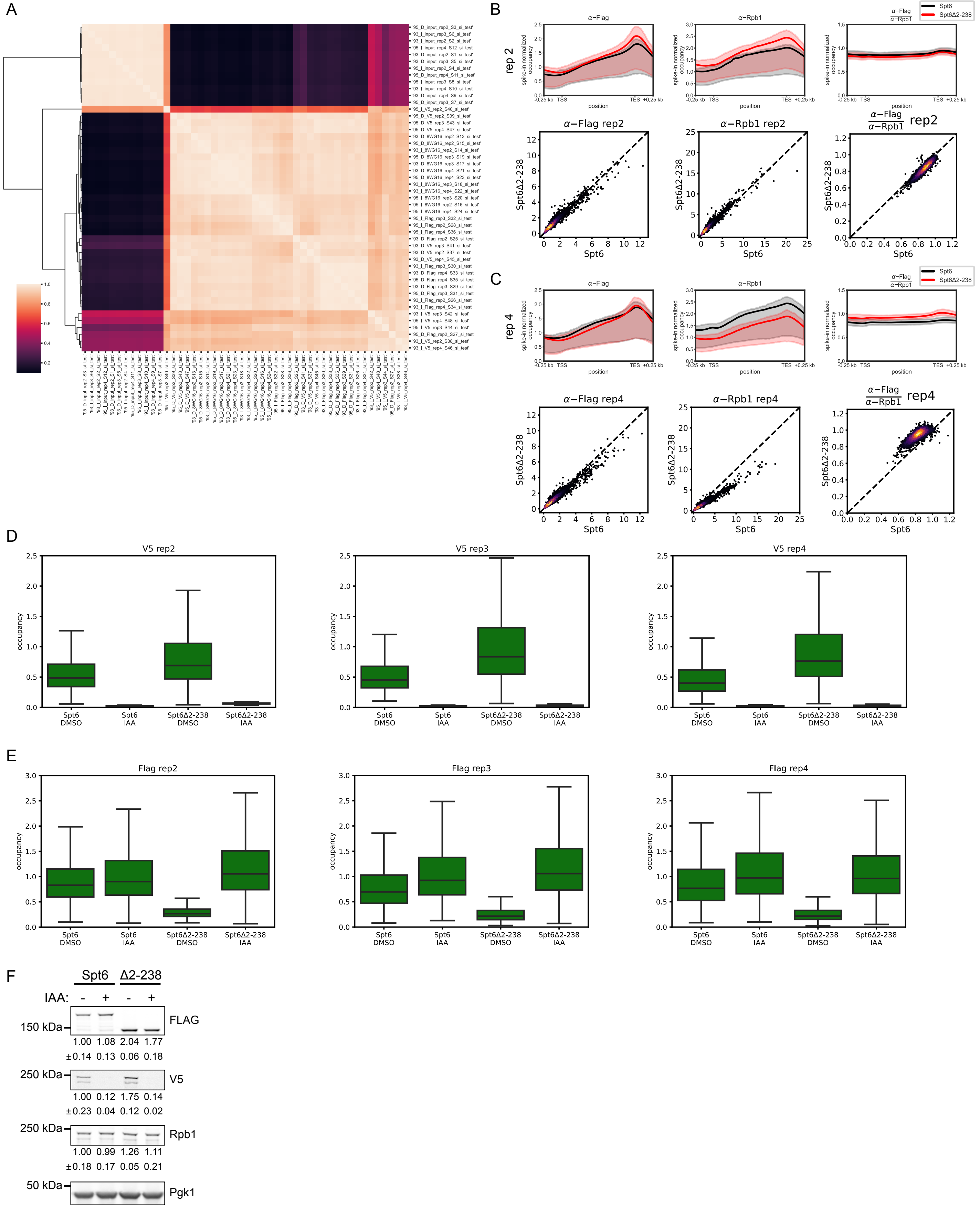

**Figure S5, related to Figure 5. Biological replicates of Spt6 and Rpb1 ChIP-seq experiments.**
(A) Heatmap of correlation between ChIP-seq biological replicates. Values are the pairwise Pearson correlation between each library of spike-in normalized ChIP-seq occupancy in all non-overlapping 200 bp bins. 93: Spt6, 95: Spt6Δ2-238, D: DMSO-treated, I: IAA-treated. Biological replicates 2, 3, and 4 are shown. Samples in replicate 1 did not sonicate well and were not used for ChIP experiments.
(B) ChIP-seq data from replicate 2. Shown above are metagene plots of spike-in normalized ChIP-seq occupancy for Flag, Rpb1, and Rpb1-normalized Flag at 3,087 non-overlapping protein-coding genes scaled to the same length. The line represents the average occupancy at each position calculated in bins of 10 bp. The shading indicates the inter-quartile range. Below are scatterplots of per-gene occupancy values for Spt6Δ2-238 cells vs. Spt6 cells. Each dot represents the average ChIP occupancy of a gene represented in the metagene. The dashed line plots $y = x$. Dots are colored using a gaussian kernel density estimate.
(C) As in (B), but for replicate 4. We observed lower Rpb1 ChIP-seq occupancy in Spt6Δ2-238 cells in this biological replicate. We did not measure a meaningful decrease in Spt6Δ2-238 association with chromatin either by Flag or Rpb1-normalized Flag ChIP-seq occupancy.
(D) Auxin treatment depletes genome-encoded Spt6 from chromatin. The mean V5 spike-in normalized ChIP-seq occupancy measurements for each of 3,087 non-overlapping protein-coding genes were plotted as a boxplot. The three biological replicates are shown side-by-side. V5 ChIP-seq occupancy is dramatically reduced in the depleted (IAA) condition compared to the non-depleted (DMSO) condition.
(E) ChIP-seq analysis of the chromatin association of Spt6Δ2-238. Boxplots as in (D), but for Flag ChIP-seq occupancy. Spt6Δ2-238 association with chromatin is substantially lower in the non-depleted (DMSO) condition than wild-type Spt6. Spt6Δ2-238 and Spt6 associate with chromatin at similar levels in the depleted (IAA) condition.
(F) Western blots showing the levels of plasmid-encoded Spt6 (Flag), genome-encoded Spt6 (V5), and Rpb1 in the samples used for ChIP-seq. Pgk1 was used as a loading control. The three biological replicates used for sequencing were analyzed by Western blot and the results from replicate 3 are shown here. The numbers below each blot indicate the abundance the measured protein in each sample relative to the wild-type Spt6 DMSO condition, normalized by Pgk1 signal. Values shown are mean ± SEM.

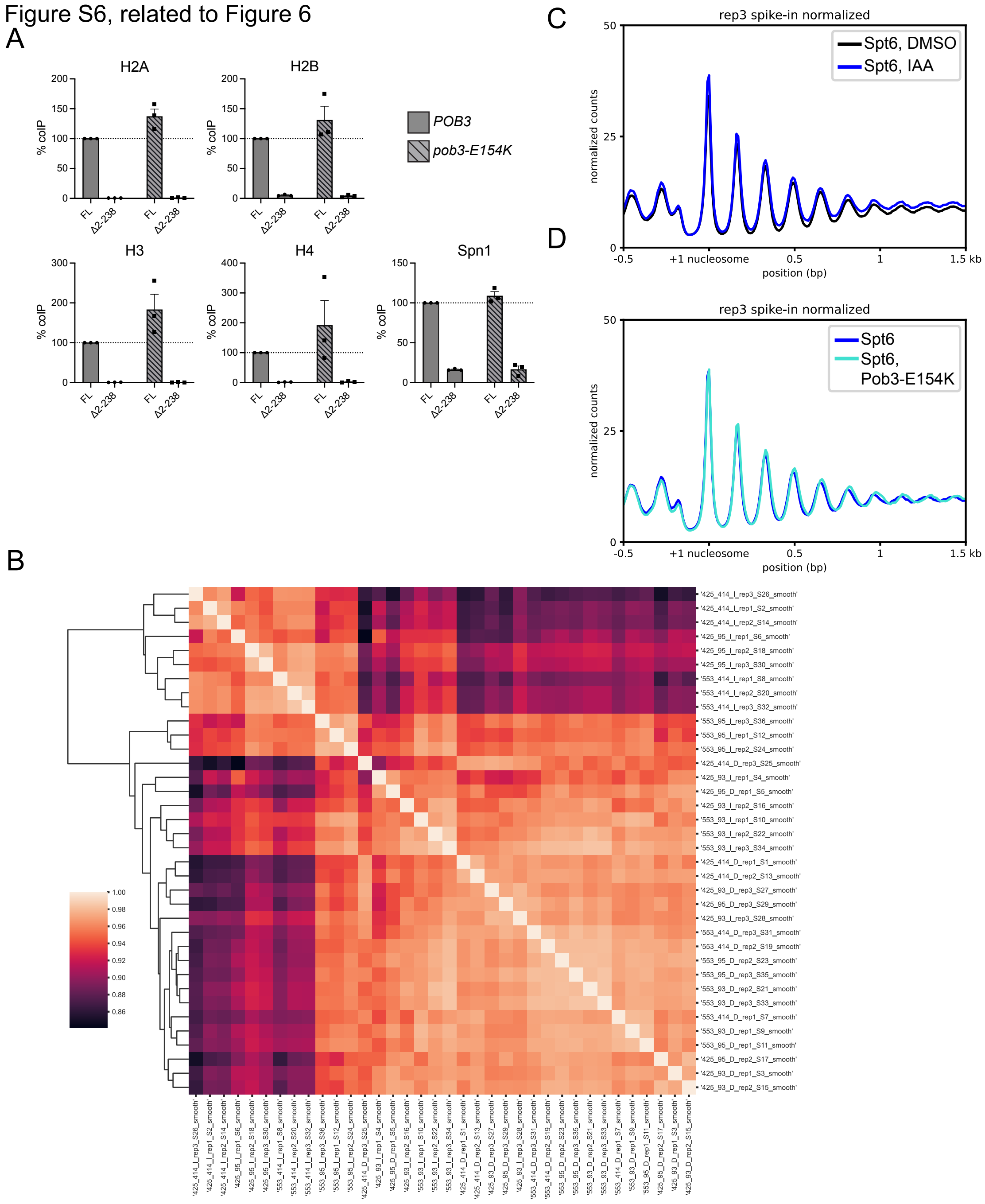


**Figure S6, related to Figure 6. The Spt6 NTD and FACT cooperate to control chromatin structure.**
(A) Quantification of coIP signal from three biological replicates of the experiment shown in Figure 6B. Values shown are mean ± SEM. FL: full-length Spt6, Δ2-238: Spt6Δ2-238.
(B) Heatmap of correlation between MNase-seq biological replicates. Values are the pairwise Pearson correlation between each library of spike-in normalized MNase-seq occupancy in all non-overlapping 75 bp bins. 425: *POB3*, 553: *pob3-E154K*, 414: empty vector, 93: Spt6, 95: Spt6Δ2-238, D: DMSO-treated, I: IAA-treated. Biological replicates 1, 2, and 3 are shown.

(C) Depletion of genome-encoded Spt6 has no effect on chromatin structure in cells expressing Spt6 from a plasmid. Metagene plot of spike-in normalized nucleosome dyad occupancy in non-overlapping 20 bp bins in cells expressing wild-type Spt6 in the presence (DMSO, black) or absence (IAA, blue) of genome-encoded Spt6. The lines represent the mean occupancy across 3,087 non-overlapping protein-coding genes. Shown is a representative of three biological replicates.
(D) The *pob3-E154K* mutation does not have an effect on chromatin structure in cells expressing wild-type Spt6. Metagene plot is as in (C).

Figure S7, related to Figure 7

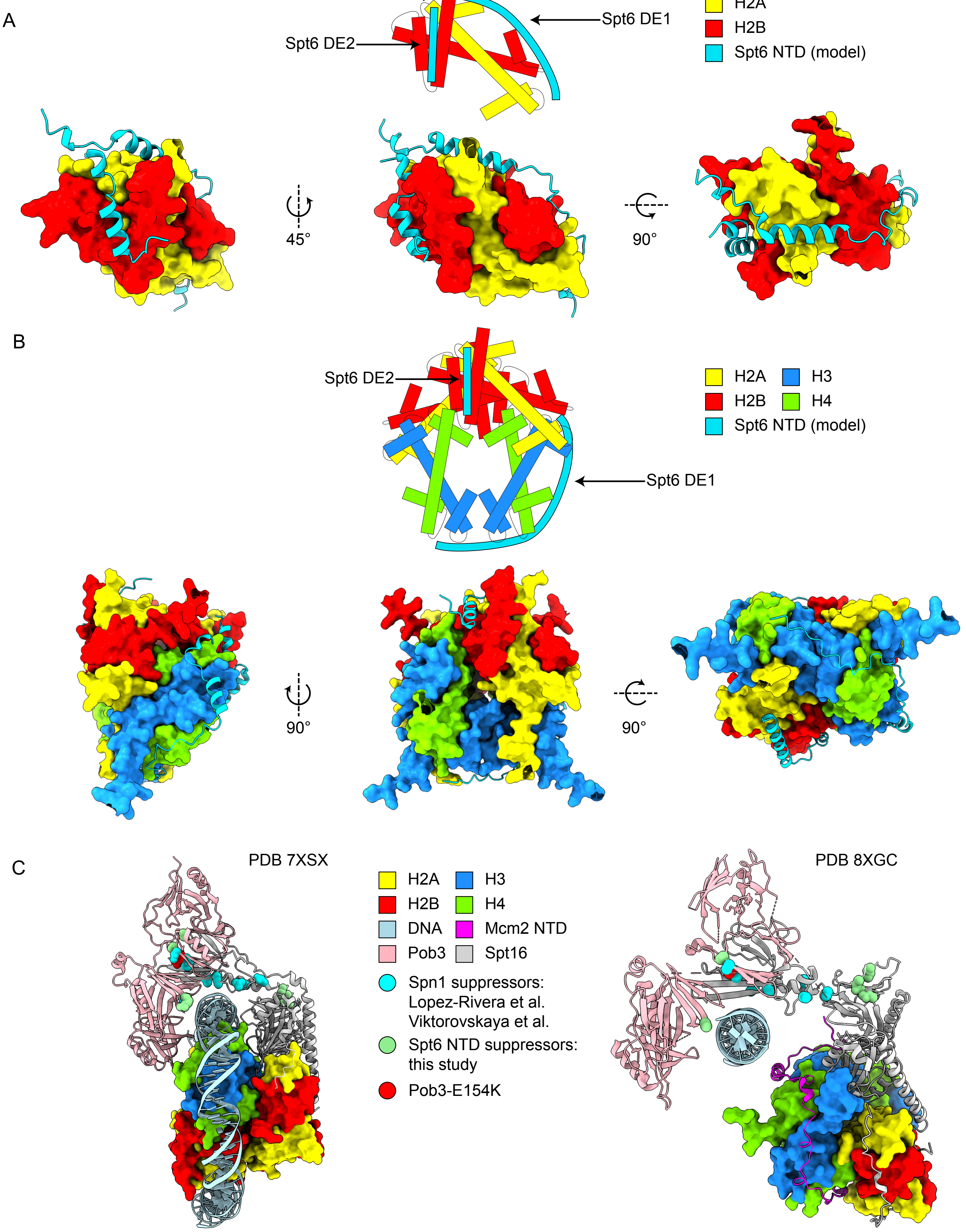

**Figure S7, related to Figure 7. Structural models supporting the cooperation between Spt6 and FACT in co-transcriptional histone transfer.**

(A) Alphafold 3 predicted structures of Spt6 NTD DE1 and DE2 interacting with a histone H2A/H2B dimer. Above is shown a cartoon representation of the predicted interactions. Below are shown three perspectives of the predicted structures. Axes and angles of rotation relative to the center structure are indicated. DE1 is predicted to interact with the DNA binding surfaces of H2A/H2B (see Figure S3), while DE2 is predicted to interact with a non-DNA binding face of histone H2B.

(B) Alphafold 3 predicted structures of Spt6 NTD DE1 and DE2 interacting with a histone octamer. Above is shown a cartoon representation of the predicted interactions. Below are shown three perspectives of the predicted structures. Axes and angles of rotation relative to the center structure are indicated. DE1 is predicted to interact with the DNA binding surface of H3/H4, while DE2 is predicted to interact with a non-DNA binding face of histone H2B (same as in (A)).

(C) Structures of FACT interacting with nucleosomes or histones and DNA. To the left is the structure of FACT bound to a nucleosome that has been partially unwrapped by transcription[3]. To the right is the structure of FACT during DNA replication in an 'open' conformation, where Pob3 and a portion of Spt16 remain associated with DNA, while the middle and C-terminal domains of Spt16 stay bound to a histone hexamer[4]. In this structure, DNA is replaced by the Mcm2 acidic NTD on the DNA binding surface of histones H3/H4. In both structures, the locations of residues mutated in FACT suppressors of Spn1-related[5,6] and Spt6 NTD-related (this study) are highlighted.

**Table S1. Phenotypes of charge substitution mutations.**

| *SPT6 allele* | Spt6 residue numbers | number of amino acids substituted | viability | 30°C | 37°C | -Lys |
|---|---|---|---|---|---|---|
| *SPT6* | | | viable | + | + | - |
| *spt6-DE1A* | 13-70 | 32 | viable | +/- | - | +/- |
| *spt6-DE2A* | 86-99 | 9 | viable | + | +/- | + |
| *spt6-DE3A* | 151-162 | 9 | viable | + | + | - |
| *spt6-DE4A* | 194-211 | 13 | viable | + | - | +/- |
| *spt6-DE5A* | 264-283 | 12 | viable | + | + | - |
| *spt6-KR1A* | 77-87 | 10 | viable | + | + | - |
| *spt6-KR2A* | 109-124 | 7 | viable | + | + | - |
| *spt6-KR3A* | 213-229 | 7 | viable | + | + | - |
| *spt6-KR4A* | 288-293 | 4 | viable | + | + | - |

Phenotypes were analyzed based on yeast growth under the indicated conditions by spot tests. Growth at 30°C and 37°C was scored after two days of incubation at the indicated temperature. Growth on SC-lysine (-Lys) was scored after three days of incubation at 30°C. Growth on -Lys indicates suppression of the *lys2-128δ* allele, the Spt- phenotype.

**Table S3. Genetic analysis of *spt6* mutations.**

| genotype | viability | 30°C | 37°C | 39°C | -Lys |
|---|---|---|---|---|---|
| analysis of *spt6 spn1* and *spt6 elf1* double mutants | | | | | |
| *spn1-F267E* | viable | + | + | +/- | +/- |
| *elf1Δ* | viable | + | + | + | - |
| *spt6ΔDE1DE2* | viable | + | + | - | + |
| *spt6ΔDE1DE2 spn1-F267E* | viable | + | +/- | - | + |
| *spt6ΔDE1DE2 elf1Δ* | inviable[1] | | | | |
| *spt6ΔDE1DE4* | viable | + | + | +/- | + |
| *spt6ΔDE1DE4 spn1-F267E* | viable | + | +/- | - | + |
| *spt6ΔDE1DE4 elf1Δ* | inviable[1] | | | | |
| *spt6ΔDE2DE4* | viable | + | - | - | + |
| *spt6ΔDE2DE4 spn1-F267E* | inviable[1] | | | | |
| *spt6ΔDE2DE4 elf1Δ* | inviable[1] | | | | |
| analysis of *spt6* and *spn1* suppressors | | | | | |
| *spt6Δ2-238* | inviable[1] | | | | |
| *pob3-E154K* | viable | + | + | + | - |
| *spt6Δ2-238 pob3-E154K* | viable | + | + | - | + |
| *spt6Δ2-238 set2Δ* | viable | -/+ | - | - | + |
| *spt6Δ2-238 chd1Δ* | viable | -/+ | - | - | + |
| *spt6Δ2-238 [SPN1 URA3]* | inviable[2] | | | | |
| *spn1Δ pob3-E154K* | viable | + | + | +/- | + |
| *spt6Δ2-238 spn1Δ pob3-E154K* | inviable[1] | | | | |

Phenotypes were analyzed based on yeast growth under the indicated conditions by replica plating. Growth at 30°C, 37°C, and 39°C was scored after two days of incubation at the indicated temperature. Growth on SC-lysine (-Lys) was scored after one day of incubation at 30°C. Growth on -Lys indicates suppression of the *lys2-128δ* allele, the Spt- phenotype.

[1]Inviable genetic combinations were determined by tetrad analysis. Strains used for tetrad analysis were as follows: FY3565 (*spt6ΔDE1DE2/SPT6 elf1Δ/ELF1*), FY3566 (*spt6ΔDE1DE4/SPT6 elf1Δ/ELF1*), FY3567 (*spt6ΔDE2DE4/SPT6 elf1Δ/ELF1*), FY3564 (*spt6ΔDE2DE4/SPT6 spn1-F267E/SPN1*), FY3577 and FY3578 (*spt6Δ2-238/SPT6 pob3-E154K/POB3*), FY3559 (*spt6Δ2-238/SPT6 spn1Δ/SPN1 pob3-E154K/pob3-E154K*). Eight to 27 tetrads were analyzed per strain, depending on the particular analysis. If the double mutant combinations were viable, we would expect to observe on average one spore with both mutations per tetrad. In all cases the spores expected to be double mutants failed to grow, indicating a synthetic lethal interaction between the mutant alleles.

[2]Failure of *SPN1* overexpression to rescue the inviability of *spt6Δ2-238* was determined by failure of strain FY3544FY3544 carrying *[spt6Δ2-238 TRP1]* and *[SPN1 URA3]* plasmids to grow in Spt6 depletion (YPD+NAA) conditions (see Methods).

**Table S4. Mutations in genes encoding FACT subunits identified in *spt6ΔDE2DE4* temperature-sensitive phenotype suppressors**

| gene | amino acid change | evidence |
|---|---|---|
| *POB3* | N141K | whole-genome sequencing |
| | A232L | Sanger sequencing |
| | R256L | Sanger sequencing |
| *SPT16* | Q666E | Sanger sequencing |
| | L669S | whole-genome sequencing |
| | N723K | Sanger sequencing |

**Table S5. Yeast strains used in this study.**

| strain name | genotype | purpose |
|---|---|---|
| FY87 | *MATα leu2Δ1 lys2-128δ ura3-52* | Parent CRISPR strain, NTD CoIPs |
| FY3276 | *MATα leu2Δ1 lys2-128δ ura3-52 SPT6-3xFLAG* | Parent CRISPR strain |
| FY37 | *MAT***a** *his3Δ200 lys2-128δ ura3-52* | Crosses genetic analysis |
| FY2377 | *MAT***a** *his4-912δ leu2Δ1 lys2-128δ ura3-52 elf1ΔNATMX* | Genetic analysis |
| FY3544 | *MATα LEU2::TIR1 lys2-128δ ura3-52 trp1Δ63 SPT6-V5-AID:KanMX* | Spt6 depletion, MNase-seq, ChIP-seq |
| FY3545 | *MATα LEU2::TIR1 lys2-128δ ura3-52 trp1Δ63 SPT6-V5-AID:KanMX spt16-L669S* | Suppressor analysis |
| FY3546 | *MAT***a** *LEU2::TIR1 lys2-128δ ura3-52 trp1Δ63 SPT6-V5-AID:KanMX pob3-E154K* | Suppressor analysis, MNase-seq |
| FY3547 | *MATα leu2Δ1 lys2-128δ ura3-52 spt6ΔDE1* | *spt6ΔDE1* |
| FY3548 | *MATα leu2Δ1 lys2-128δ ura3-52 spt6ΔDE2* | *spt6ΔDE2* |
| FY3549 | *MATα leu2Δ1 lys2-128δ ura3-52 spt6ΔDE4* | *spt6ΔDE4* |
| FY3550 | *MATα leu2Δ1 lys2-128δ ura3-52 spt6ΔDE1DE2* | *spt6ΔDE1DE2* |
| FY3551 | *MATα leu2Δ1 lys2-128δ ura3-52 spt6ΔDE1DE4* | *spt6ΔDE1DE4* |
| FY3552 | *MATα leu2Δ1 lys2-128δ ura3-52 spt6ΔDE2DE4* | *spt6ΔDE2DE4*, suppressor selection |
| FY3553 | *MATα leu2Δ1 lys2-128δ ura3-52 spt6ΔDE2DE4 pob3-N141K* | initial *pob3*-sup |
| FY3554 | *MATα leu2Δ1 lys2-128δ ura3-52 spt6ΔDE2DE4 spt16-L669S* | initial *spt16*-sup |
| FY3555 | *MAT***a** *his3Δ200 leu2Δ1 lys2-128δ ura3-52 spt6ΔDE2DE4* | Testing suppressors for dominance |
| FY3556 | *MATα leu2Δ1 lys2-128δ ura3-52 spn1-F267E* | *spn1-F267E* |
| FY3557 | *MAT***a** *his3Δ200 lys2-128δ ura3-52 pob3-E154K* | Suppressor analysis |
| FY3558 | *MATα leu2Δ1 lys2-128δ ura3-52 spt6Δ2-238 pob3-E154K* | Suppressor analysis |
| FY3559 | *MAT***a***/MATα his3Δ200/HIS3 leu2Δ1/LEU2 lys2-128δ/lys2-128δ ura3-52/ura3-52 pob3-E154K/pob3-E154K spt6Δ2-238/SPT6 spn1ΔKanMX/SPN1* | Suppressor analysis |
| FY3560 | *MAT***a** *leu2Δ1 lys2-128δ ura3-52 trp1Δ63 spt6Δ2-238 chd1ΔKanMX* | Suppressor analysis |
| FY3561 | *MAT***a** *his3Δ200 leu2Δ1 lys2-128δ ura3-52 spt6Δ2-238 set2ΔNatMX* | Suppressor analysis |
| FY3562 | *MAT***a***/MATα his3Δ200/HIS3 leu2Δ1/LEU2 lys2-128δ/lys2-128δ ura3-52/ura3-52 spn1-F267E/SPN1 spt6ΔDE1DE2/SPT6* | Suppressor analysis |
| FY3563 | *MAT***a***/MATα his3Δ200/HIS3 leu2Δ1/LEU2 lys2-128δ/lys2-128δ ura3-52/ura3-52 spn1-F267E/SPN1 spt6ΔDE1DE4/SPT6* | Suppressor analysis |
| FY3564 | *MAT***a***/MATα his3Δ200/HIS3 leu2Δ1/LEU2 lys2-128δ/lys2-128δ ura3-52/ura3-52 spn1-F267E/SPN1 spt6ΔDE2DE4/SPT6* | Suppressor analysis |

| | | |
|---|---|---|
| FY3565 | MAT**a**/MATα his4-912δ/HIS4 leu2Δ1/leu2Δ1 lys2-128δ/lys2-128δ ura3-52/ura3-52 elf1ΔNatMX/ELF1 spt6ΔDE1DE2/SPT6 | Suppressor analysis |
| FY3566 | MAT**a**/MATα his4-912δ/HIS4 leu2Δ1/leu2Δ1 lys2-128δ/lys2-128δ ura3-52/ura3-52 elf1ΔNatMX/ELF1 spt6ΔDE1DE4/SPT6 | Suppressor analysis |
| FY3567 | MAT**a**/MATα his4-912δ/HIS4 leu2Δ1/leu2Δ1 lys2-128δ/lys2-128δ ura3-52/ura3-52 elf1ΔNatMX/ELF1 spt6ΔDE2DE4/SPT6 | Suppressor analysis |
| FY3568 | MATα leu2Δ1 lys2-128δ ura3-52 spt6-DE1A | spt6-DE1A |
| FY3569 | MAT**a** his3Δ200 lys2-128δ ura3-52 spt6-DE2A | spt6-DE2A |
| FY3570 | MATα his3Δ200 leu2Δ1 lys2-128δ ura3-52 spt6-DE3A | spt6-DE3A |
| FY3571 | MATα leu2Δ1 lys2-128δ ura3-52 spt6-DE4A | spt6-DE4A |
| FY3572 | MATα his3Δ200 leu2Δ1 lys2-128δ ura3-52 spt6-DE5A | spt6-DE5A |
| FY3573 | MAT**a** leu2Δ1 lys2-128δ ura3-52 spt6-KR1A | spt6-KR1A |
| FY3574 | MATα his3Δ200 leu2Δ1 lys2-128δ ura3-52 spt6-KR2A | spt6-KR2A |
| FY3575 | MAT**a** leu2Δ1 lys2-128δ ura3-52 spt6-KR3A | spt6-KR3A |
| FY3576 | MATα leu2Δ1 lys2-128δ ura3-52 spt6-KR4A | spt6-KR4A |
| FY3577 | MAT**a**/MATα his3Δ200/HIS3 leu2Δ1/LEU2 lys2-128δ/lys2-128δ ura3-52/ura3-52 pob3-E154K/POB3 spt6Δ2-238/SPT6 | Suppressor analysis |
| FY3578 | MAT**a**/MATα his3Δ200/HIS3 leu2Δ1/LEU2 lys2-128δ/lys2-128δ ura3-52/ura3-52 pob3-E154K/POB3 spt6Δ2-238/SPT6 | Suppressor analysis |

**Table S6. Plasmids used in this study.**

| name | contains | source | purpose |
|---|---|---|---|
| pRS414 | pRS414 [TRP1, CEN/ARS] | PMID: 2659436 | Cloning, spot tests, MNase-seq |
| pJLW93 | pRS414 [SPT6-3xFLAG, TRP1, CEN/ARS] | This study | Spot tests, MNase-seq |
| pJLW95 | pRS414 [spt6Δ2-238-3xFLAG, TRP1, CEN/ARS] | This study | Spot tests, MNase-seq |
| pJLW158 | pRS414 [spt6ΔDE1DE2DE4-3xFLAG, TRP1, CEN/ARS | This study | Spot tests |
| GHB346 | pRS316 [SPN1, URA3, CEN/ARS] | Grant Hartzog | spt6 suppressor analysis |
| pML104 | Cas9, sgRNA, URA3, 2µ | PMID: 26305040 | CRISPR genome editing |
| pFA6a-KanMX6 | pFA6 [KanMX] | PMID: 9717240 | Strain construction |
| pJLW112 | 6xHisMBP-3C-Spt6(1-335)-Cys-3xFLAG expression vector | This study | Protein expression |
| pJLW123 | 6xHisMBP-3C-Spt6(1-238)-Cys-3xFLAG expression vector | This study | Protein expression |
| pJLW72 | 6xHis-H2A/H2B coexpression in pET-LIC vector (*S. cerevisiae*) | This study | Protein expression |
| p416-GAL1 | pRS416 [pGAL1:, URA3, CEN/ARS] | PMID: 2659436 | Cloning, coIP |
| pJLW140 | pRS416 [pGAL1:Spt6(1-335)-Cys-3xFLAG, URA3, CEN/ARS] | This study | coIP |
| pJLW141 | pRS416 [pGAL1:Spt6(1-238)-Cys-3xFLAG, URA3, CEN/ARS] | This study | coIP |
| pJLW142 | pRS416 [pGAL1:Spt6(239-335)-Cys-3xFLAG, URA3, CEN/ARS] | This study | coIP |
| pJLW156 | pRS416 [pGAL1:Spt6(1-335)ΔDE1DE2DE4-Cys-3xFLAG, URA3, CEN/ARS] | This study | coIP |
| pJLW159 | pRS416 [pGAL1:Spt6(1-335)ΔDE1DE2-Cys-3xFLAG, URA3, CEN/ARS] | This study | coIP |
| pJLW160 | pRS416 [pGAL1:Spt6(1-335)ΔDE1DE4-Cys-3xFLAG, URA3, CEN/ARS] | This study | coIP |
| pJLW161 | pRS416 [pGAL1:Spt6(1-335)ΔDE2DE4-Cys-3xFLAG, URA3, CEN/ARS] | This study | coIP |
| pJLW165 | pRS416 [pGAL1:SUPT6H(1-305)-Cys-3xFLAG, URA3, CEN/ARS] | This study | coIP |
| pJLW166 | pRS416 [pGAL1:SUPT6H(1-206)-Cys-3xFLAG, URA3, CEN/ARS] | This study | coIP |

| | | | |
|---|---|---|---|
| Spn1-pMCSG7 | His6, TEV cleavage site, Spn1(1-410) | This study | Protein expression (NMR) |
| Spn1(1-410)-pMCSG7 | His6, TEV cleavage site, Spn1(149-410) | This study | Protein expression (NMR) |
| Spn1(149-410)-pMCSG7 | His6, TEV cleavage site, Spn1(149-296) | This study | Protein expression (NMR) |
| Elf1 -pMCSG7 | His6, TEV cleavage site, Elf1 (1-145) | This study | Protein expression (NMR) |
| IWS1(550-692)-pMCSG7 | His6, TEV cleavage site, IWS1(550-692) | Prepared in Veverka lab PMID: 34822292 | Protein expression (NMR) |
| Spt6(1-206)-pMCSG7 | His6, GB1, TEV cleavage site, hSpt6(1-206) | This study | Protein expression (NMR) |
| Spt6(1-248)-pMCSG7 | His6, GB1, TEV cleavage site, hSpt6(1-248) | Prepared in Veverka lab PMID: 34822292 | Protein expression (NMR) |
| H2A-pMCSG7 | histone H2A (*X. laevis*) | This study | Protein expression (NMR) |
| H2B-pMCSG7 | histone H2B (*X. laevis*) | This study | Protein expression (NMR) |
| H3-pMCSG7 | histone H3 (*X. laevis*) | This study | Protein expression (NMR) |
| H4-pMCSG7 | histone H4 (*X. laevis*) | This study | Protein expression (NMR) |
